# Crowder induced conformational fluctuations modulate the phase separation of yeast SUP35 NM domain

**DOI:** 10.1101/2024.08.28.610075

**Authors:** Sumangal Roychowdhury, Sneha Menon, Narattam Mandal, Jagannath Mondal, Krishnananda Chattopadhyay

**Affiliations:** Protein Folding and Dynamics Laboratory, Structural Biology and Bioinformatics Division, CSIR-Indian Institute of Chemical Biology 4, Raja SC Mullick Road, Kolkata 700032, India; Tata Institute of Fundamental Research Hyderabad, 36/P, Gopanapally village, Serilingampally mandal, Hyderabad 500046, India

**Keywords:** Liquid-liquid phase separation, prion, yeast, fluorescence, biophysics, spectroscopy, Simulation

## Abstract

Intrinsically disordered proteins (IDPs) like Sup35NM can undergo liquid-liquid phase separation (LLPS) to form biomolecular condensates, a process influenced by their conformational flexibility and the crowded intracellular environment. This study investigates how molecular crowding, specifically the size and shape of crowders like Dextran and Ficoll, modulates the conformational states and phase separation behavior of Sup35NM. Using fluorescence correlation spectroscopy (FCS) and molecular dynamics simulations, we observed that Dextran, depending on its molecular weight, induces both compaction and expansion of Sup35NM, driving phase separation at certain thresholds. Notably, rod-like Dextran crowders promote phase separation, while spherical Ficoll does not, highlighting the impact of crowder geometry on IDP behavior. Computational modelling further revealed that the crowder shape influences Sup35NM’s conformational ensemble by modulating intra- and inter-domain interactions. These findings elucidate the role of crowding agents in IDP phase behavior, suggesting that cellular crowding may regulate IDP functionality through conformational control.

## Introduction

Intrinsically disordered proteins (IDPs) can assume multiple conformational states depending on their environment and localization. This structural flexibility of IDPs can significantly influence their activities and involvement in multiple processes, including intracellular signalling, active protein transport, metabolic processes, and protein aggregation^1^. Recent advances in biophysical chemistry have revealed the ability of IDPs to form biomolecular condensates through homotypic and heterotypic interactions they encounter in the solution ^2^. There is accumulating evidence that this process of liquid-liquid phase separation (LLPS) may be critical in the formation of membraneless organelles, which play key roles in cellular organization and regulation of intracellular biochemical processes even in transcription and translation machinery^3^. While *in-vitro* characterization of phase separation has mostly been carried out under dilute solution conditions, investigations on the role of molecular crowding on phase separation are now being pursued ^4^. It is established that molecular crowding owing to the excluded volume effect can alter the phase separation capability of IDPs by modulating the intra- and inter-molecular interactions. To mimic the cytoplasmic environment in vitro, inert molecules such as ficoll, dextran, polyethylene glycol, and protein molecules (like serum albumin and lysozyme) have been routinely used as crowding agents^5^. It has been demonstrated that both the molecular weight and hydrodynamic size of the crowding agents and the nature of probe molecules play crucial roles in the behaviour of a crowder^6^.

In yeast, the protein Sup35 plays an essential role in translation termination when functioning in conjunction with another protein, Sup45. Sup35 comprises three domains: the N-terminal domain (aa: 1-74), the Middle domain (M domain, aa: 75-253), and the C-terminal domain (aa: 254-685)^7^. It has been shown recently that in response to transient stress, intrinsically disordered N-terminal and middle regions (NM) of Sup35 (together a 253 amino-acid long sequence, Sup35NM, (Figure 1a & S1a), confer stress-sensing abilities and facilitate the phase separation into biomolecular condensates^8^. Intriguingly, the same NM domain is also required for Sup35 to misfold and aggregate in association with the [PSI+] prion^7^.

**Figure 1:**
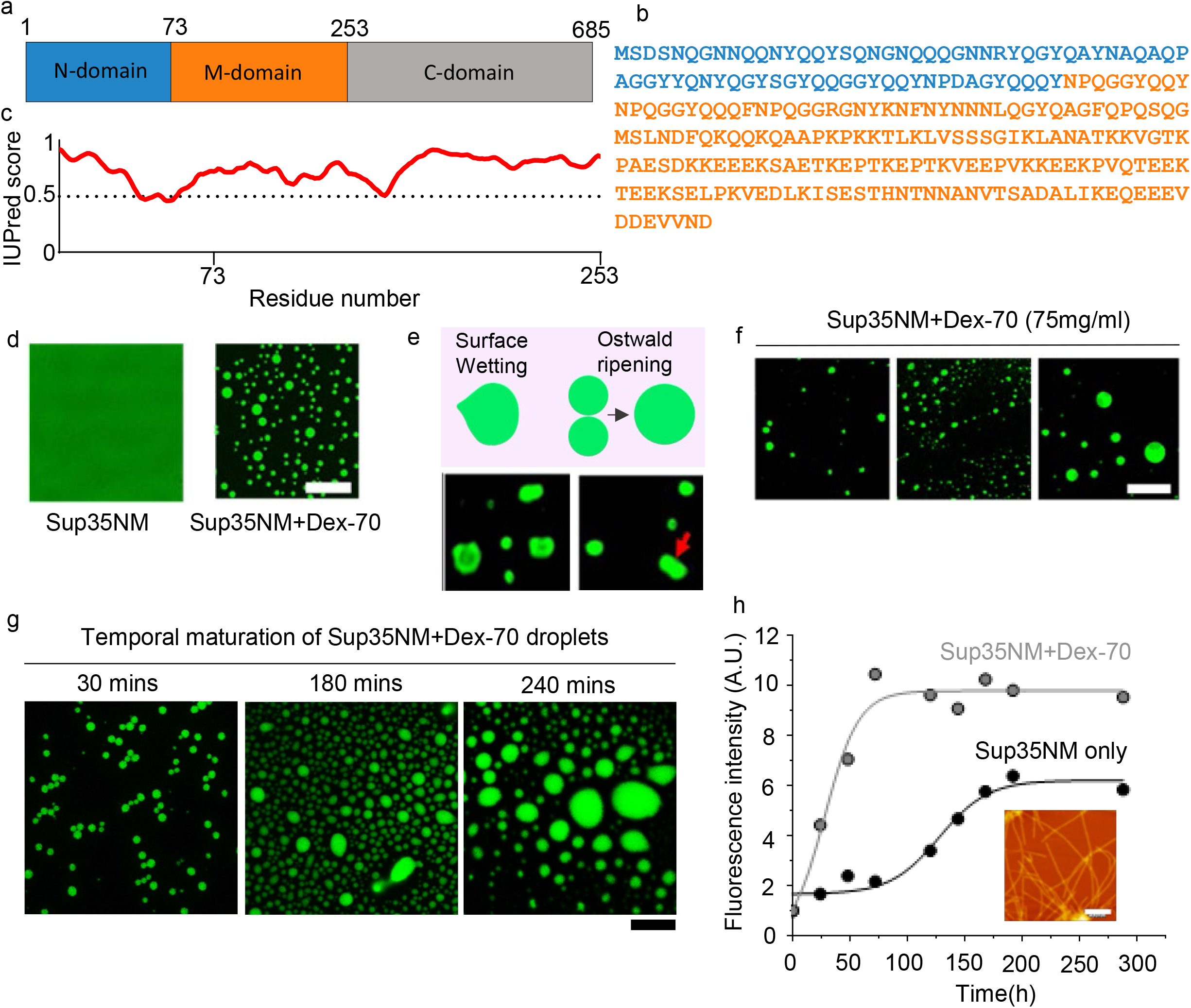
Phase separation of Sup35NM in the presence of Dextran: (a) The schematic diagram of the full-length Sup35 protein indicating different domains. (b) The amino acid sequence of full-length Sup35 protein. (c) The disorder prediction of Sup35 from linear sequence analysis using IUPred2 prediction. The scores above the dashed line (0.5) suggest the disorder in the sequence. (d) Confocal microscopic image using 10µM Sup35NM (for all images 50nM Rh-green labelled SUP35 was mixed with unlabelled proteins) in the absence of Dextran-70 (left), no droplet formation was observed along with the representative confocal image of liquid droplets formed in the presence of 75 mg/ml dextran-70 (right); Scale bar: 10µM (e) Fluorescent images of Sup35 liquid droplets showing typical liquid-like behavior of Oswald ripening and surface wetting. (g) Representative confocal images during temporal maturation of liquid droplets over 240 mins. Scale bar: 10µM. (h) A plot of fluorescence intensity vs. time for Thioflavin-T (ThT) binding to show aggregation of Sup35NM in the presence of Dextran-70 100 mg/ml. Representative AFM image of amyloid fibrils formed in the presence of 100 mg/ml dextran-70. Scale bar: 400nm.

The conformational plasticity of an IDP allows the protein to sample through multiple conformations. The population distribution of the conformational states of an IDP would vary based on multiple factors, including solution environments, pH and the presence of excluded volume effects^9^. Though the properties of the phase transition of an IDP have been shown to depend on the molecular weight and the shape of a crowder, the effect of the nature of a crowder on the population dynamics and phase separation behaviour of an IDP is still an untouched area of exploration. In this study, we used a combination of fluorescence correlation spectroscopy (FCS) and computational model to investigate the conformation-phase separation landscape of Sup35NM to find out how the size and shape of an inert crowder modulate the protein’s landscape. In a previous study, we showed that synthetic crowders affect the populations of a protein’s extended and compact states by modulating its conformational stability^10^. In the present study, which used Dextran and Ficoll as the model crowders, we show that the population distribution of the extended and compact states depends on multiple factors, including the concentrations of the protein as well as the crowders, the molecular weight and the nature of the crowder. It may be noted that Dextran is a flexible rod-like polymer of glucose, while Ficoll is relatively spherical, and hence, a comparison between Dextran 70 and Ficoll 70 (they would have similar molecular masses) would provide an understanding of the crowding effect of crowder molecule of different shape^11^. We also used different variants of dextran according to their molecular weights to illuminate the effect of size, shape and molecular weight of crowder molecule on conformational fluctuation induced LLPS propensity of SUP35 protein.

## Results and Discussion

### Sup35-NM forms biomolecular condensates in the presence of Dextran

Physicochemical properties of our model protein SUP35NM have been assessed using a number of computational and bioinformatics tools, for example, using IUPred2 and NPCR plots, we determined the charge distribution and disorderedness throughout the primary sequence of the protein^12,13^. Using IUPred2, we found that the NM domain of Sup35 is significantly disordered (Figure 1c). While Sup35NM carries a net negative charge of -5.5 at physiological pH but the M domain contains clusters of highly charged residues (Figure S1b). Prion-like Amino Acid Composition (PLAAC) tool suggests that the region of 1-141 contains prion-like low complexity sequence (Figure S1c). In addition, analysis using Predictor of Natural Disordered Regions (PONDR) shows the presence of several segments of naturally disordered regions, generally responsible for multivalent as well as chain-solvent interactions^14^ (Figure S1d). For the phase separation propensity, we used sequence-dependent predictor, FuzDrop, which predicts the residue-specific propensity score (P_DP_)^15^. The FuzDrop score was high for the entire NM domain (Figure S1e) and when we compared Sup35 with other prototypical IDPs, like alpha-Synuclein (*α*-syn) and different isoforms of Tau, using another computational tool catGRANULE, we found higher propensity of Sup35NM towards phase separation compared to the other two IDPs ^16^(Figure S1f).

It has been widely reported that synthetic water soluble polymeric crowders like polyethylene glycol (PEG), dextran, ficoll etc., can induce biomolecular condensate formation of proteins, nucleic acids through various associative, non-associative interactions and altering their conformational dynamics ^17^. At first, we studied the phase separation behavior of SUP35NM in presence of different concentrations (varying between 0 mg/ml and 100 mg/ml) of dextran 70 which is an inert rod like short branched polymer molecule. To elucidate the effect of molecular weight and shape of the crowder onto protein conformation we used other dextran variants and their arrangement in molecular weight and radius of dextran is as follows dextran 70>dextran 40>dextran 20>dextran 6^18^. For all sets of imaging experiments, we used 50nM labelled protein as a probe for imaging the droplets. Different concentration of unlabelled protein was added in the solution to figure out the ideal experimental condition of SUP35NM for further investigation of protein behavior. The initial solution condition was 10µM unlabelled protein in presence of 75mg/ml dextran 70 which produced distinctive protein droplets for further characterization. We incubated all samples identically at 370C for ∼ 30 minutes and under 180 rpm shaking conditions. We found out that the protein by itself did not phase-separate at experimental concentrations (Figure 1d, Left) but beyond a threshold concentration of Dextran-70, Sup35NM formed spherical liquid droplets (Figure 1d, Right). The droplets showed ripening and wetting, which are typical characteristics of protein droplets19 (Figure 1e) with respect to change of time. We also encountered that this phase separation behavior and dynamic properties of condensates changes with respect to both protein and crowder mass concentration. However, at higher protein concentrations (20μM in Figure 1f), the number of droplets decreased but their size increased considerably, presumably because of Ostwald ripening. It was observed earlier that liquid droplets can mature into viscoelastic liquid or solid-like aggregates based on the number and position of sticker and spacer residues in the sequence20. To study the temporal maturation of the droplets, we used identical concentrations of Sup35NM (in the presence of 75 mg/ml dextran 70), which we incubated at 37°C for different time intervals. It took approximately half an hour for the droplets to be microscopically visible. The number of droplets initially went up with time and then started to decrease after certain incubation time period as the droplet size grew bigger with significant loss of sphericity following maturation phase of condensates (Figure 1g & S1g). When we incubated the proteins for even longer duration, we observed formation of amyloid fibril-like aggregates which increased thioflavin T fluorescence with a mechanism similar to nucleation-condensation-conversion (Figure 1h). The fibrillar nature of the aggregates was confirmed by Atomic Force Microscopy (AFM) (Figure 1h, inset). As determined from the AFM data, the typical width of the fibrils was 2.7nm and ∼ 52nm, while their height was found to be ∼ 4.77 nm (Figure S1h). It may be noted that amyloid fibrils of similar characteristics were found in earlier reports for Sup35NM^21^.

### Sup35NM Phase Separation depends on the size and concentration of the crowder molecules

To obtain further insights into the effect of crowder molecule on phase separation of Sup35NM, we systematically studied the effect of different dextran variants and sup35NM concentrations in the presence of Dextran 20, Dextran 40 and Dextran 70 (Figure 2). The hydrodynamic radii of dextran 70, dextran 40 and dextran 20 are reported to be ∼6.8 nm, 4.8 nm and ∼3.24 nm respectively^22,23^. We found that the protein phase separates in presence of all three dextran variants but at distinct protein and crowder concentration, even the phase separation behaviour of SUP35 changes with respect to different variants of dextran molecules. The size of the protein droplets increased as we increased the concentration of both protein and crowder molecule highlighting the associative interaction of proteins due to excluded volume effect, however for a fixed concentration of protein and Dextran, the size of the droplets did not follow a general trend respect to the molecular weight of the Dextran (Dextran 20 vs. Dextran 40 vs. Dextran 70). For example, 62.5 mg/ml Dextran 20 and Dextran 40 produced biomolecular condensate at protein concentration of 5µM whereas no droplet was observed for the identical concentration of Dextran 70. In contrast, for 20 μM protein 50mg/ml of Dextran 20 and Dextran-40 has shown to initiate the droplet formation, while for Dextran 70, we needed lesser (37.5 mg/ml) concentration for the droplet formation. This less dextran -70 concentration requirement owing to the increased soft interaction between protein molecules due to high branching and viscosity of dextran-70 solution^22^. The scattering intensity data (optical turbidity at 600nm) also followed the trend as observed in the confocal microscopy experiment (Figure S2a). We found that the circularity increased within the phase separation regime from low crowder concentration (50 mg/ml) to high crowder concentration (100 mg/ml) for Dextran-70 and similar trends were also observed for Dextran-40 and Dextran-20. Interestingly, the deviation of circularity from a value of one (Circularity of one essentially means a perfectly circular particle) was observed for Dextran-70 was followed by Dextran-40 and Dextran-20 (Figure S2b). The qualitative and quantitative changes of phase separated protein droplets may be due to the molecular weight, branching and shape of the dextran molecules.

**Figure 2:**
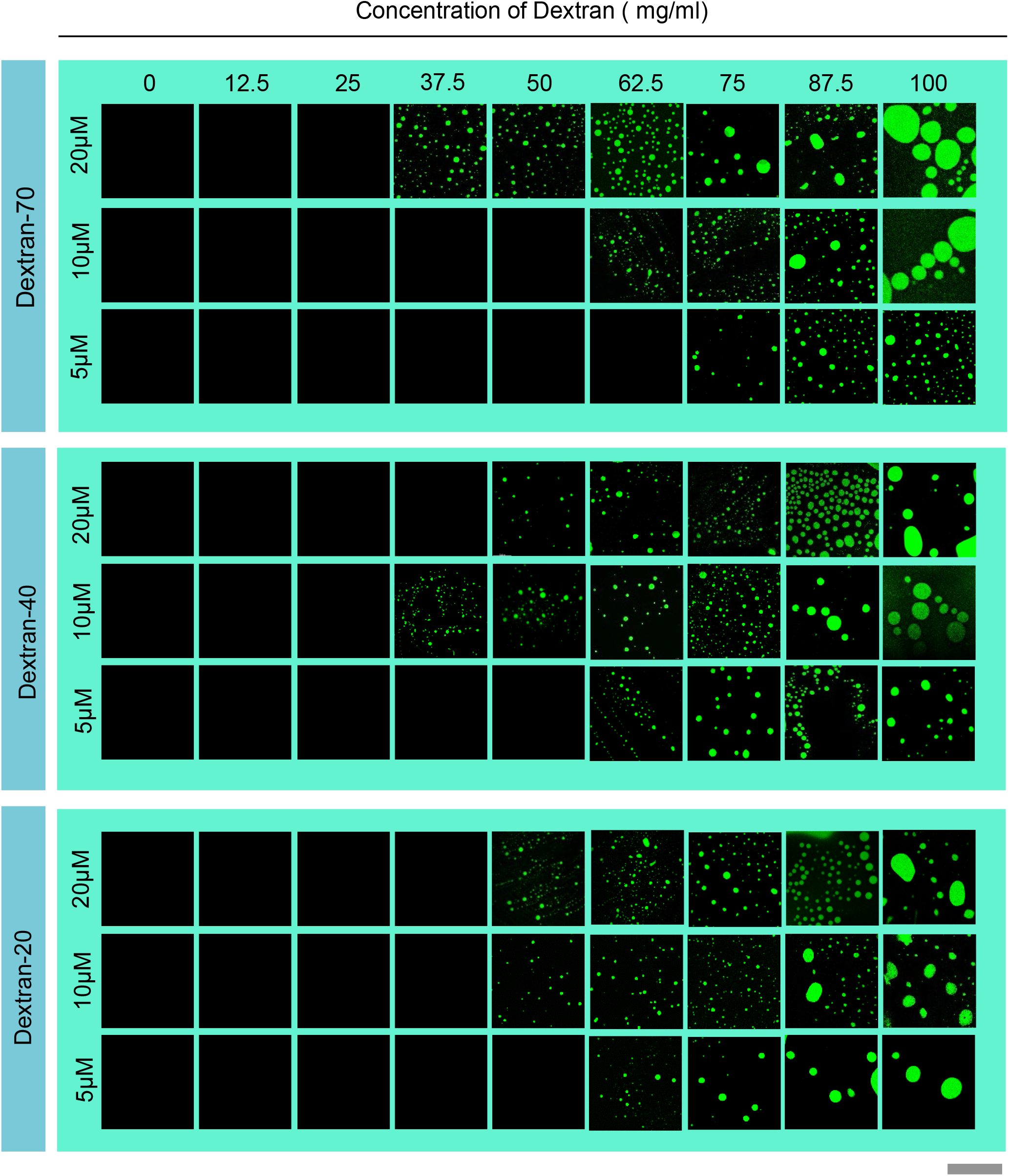
Phase diagram of Sup35NM in the presence of different crowder: Phase diagram of Sup35 in the presence of different types of Dextrans incubated for ∼30 minutes at 180 rpm shaking condition. The protein final concentrations are indicated in the left y-axis and dextran concentrations are indicated in the top. Scale bar: 20µm.

In order to investigate if the shape of the crowding agents can modulate the phase separation of Sup35NM, we then used spherical crowder molecule Ficoll-70. It may be re-emphasized that Dextran 70 is rod-like (Dextran), while Ficoll-70 is spherical (Ficoll), although both have similar molecular weight. Keeping other experimental conditions for Ficoll-70 identical to those used for Dextran experiments (as above), we found that for all protein and Ficoll-70 concentrations, there was no droplet formation for Sup35NM (Figure S2c).

### Sup35 exhibits compact (C), intermediate (N) and extended (U) conformation in different concentration regimes of the crowder

An inert crowder molecule, like Dextran, does not typically interact directly with a protein^18^. ^1^H NMR spectra of protein in the absence and presence of Dextran 70 remained superimposable, suggesting that this is likely to be true for Sup35NM as well (Figure S3). Since SUP35NM contains significant disordered region, and IDPs have been shown to sample multiple conformational states. To get an insight into the conformational heterogeneity of Sup35NM we ran a large number of FCS experiments (typically 100 with short 10 sec) at different conditions. FCS is a fluorescence-based technique that monitors fluorescence fluctuations of diffusing molecules arising in and out of a small confocal volume. FCS uses very low concentration (typically in the ∼nM concentration) of the probe, which rules out the unwanted contributions from aggregates etc^24^. In the absence of any crowder (50nM labeled Sup35NM in 20mM sodium phosphate buffer, pH 7.4), we observed a broad distribution of diffusion coefficient (D) values, which is expected for a protein with significant conformational heterogeneity (Figure S4a-c). From the distribution, we obtained a mean hydrodynamic radius (r_Hm_) of 18.5 Å that is consistent with our previously reported work with Sup35NM^21^. In the presence of 50 mg/ml Dextran70 concentration, we found that the distribution of D became narrower and shifted to the lower range of hydrodynamic radius (Figure S4a-c) whereas in high crowder concentration (100mg/ml) the distribution shifted to higher range of r_Hm_ value (Figure S4a-c). When we change the crowder molecule to Dextran 40 and 20 we found that the similar phenomenon happening but in different protein and crowder concentration (Figure S4a-c). We believe that this conformational heterogeneity of the protein is due to compaction and expansion of protein chain (Figure 3a), which populate preferentially in lower and higher range of crowder concentrations respectively. We also studied systematically Sup35NM concentrations dependence of the above equilibria (which is shown in Figure 3a) using FCS. For these FCS experiments, we used 50nM labelled proteins in the presence of different concentrations of unlabelled Sup35NM (Condition-I to Condition-IV as shown in Figure 3b).

**Figure 3:**
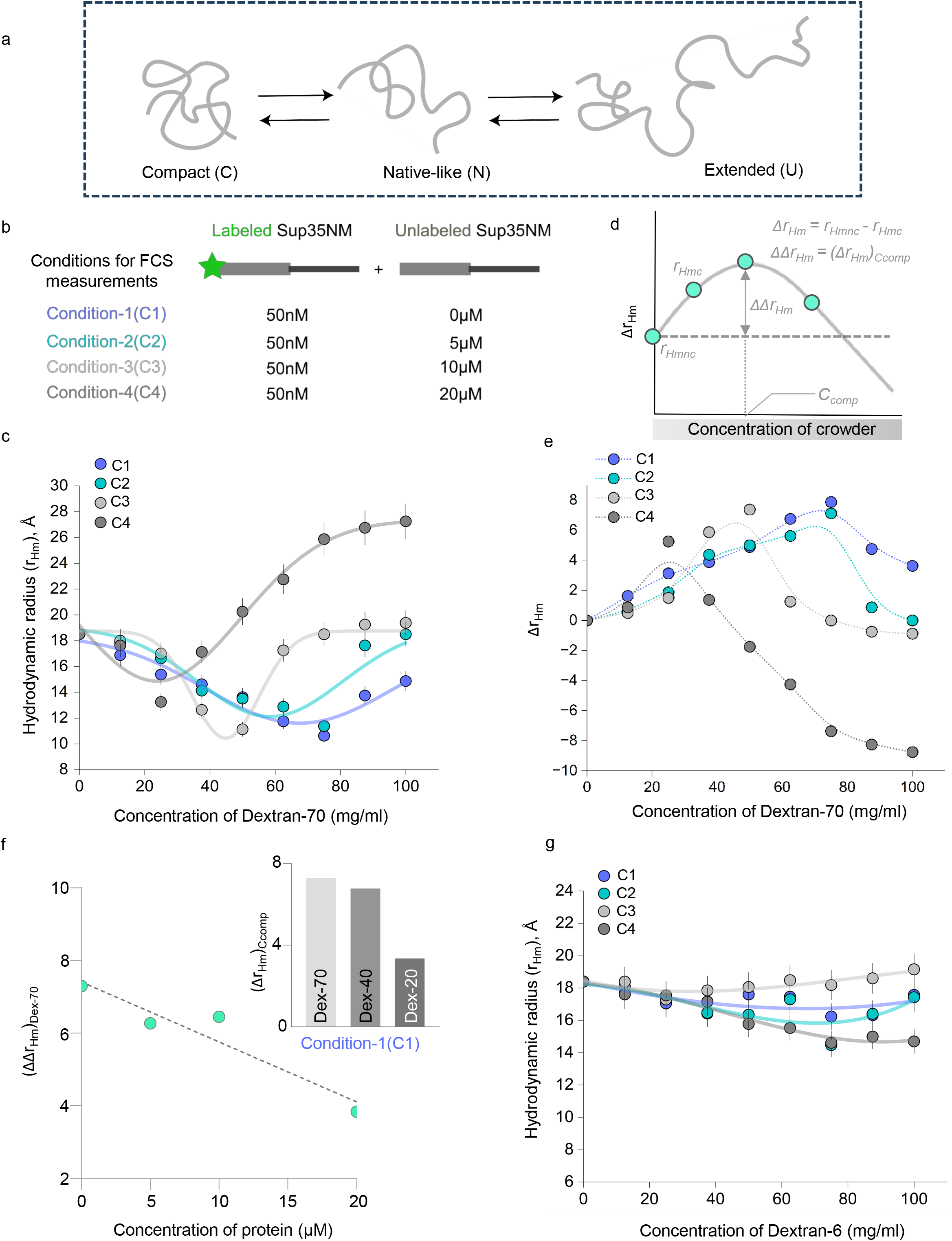
Conformational behaviour of Sup35NM under crowded conditions: (a) Schematic representation of equilibrium between different conformational states of Sup35NM domain. (b) Experimental conditions used for FCS experiments for different crowding agents. (c) Plot of hydrodynamic radius (r_Hm_), Angstrom vs. concentration (mg/ml) of crowder for Dextran-70. Purple circles indicate the 50nM Sup35NM, Cyan circles indicate 50nM labeled protein in 5µM unlabeled protein, Light grey circles indicate 50nM labeled protein in 10µM unlabeled protein, and Deep grey circles indicate 50nM labeled protein in 20µM unlabeled protein. All measurements were take n=3. The data are shown mean ± SD. (d) Schematic figure for the calculation of extent of compaction (Δr_Hm_). Δr_Hm_ is the difference between r_Hmnc_ and r_Hm_c, where r_Hmnc_ is the value of r_Hm_ in the absence of any crowding agent and r_Hmc_ is the value of r_Hm_ at any crowder concentration. C_comp_ is the concentration of the crowder at which the compaction is maximum. (ΔΔr_Hm_) is the value of (Δr_Hm_)_Ccomp_. (e) Plot of (Δr_Hm_) at different crowder concentrations for Dextran-70 at varying protein concentration (C1-C4). (f) Plot of (ΔΔr_Hm_) vs. protein concentration indicated in the x-axis. Inset : Plot of (Δr_Hm_)_Ccomp_ for different crowders (Dex-70, Dex-40, Dex-20) at condition C1. (e) Plot of hydrodynamic radius (r_H_) vs. concentration of crowder for Dextran-6. Purple circles indicate the 50nM Sup35, Cyan circles indicate 50nM labeled protein in 5µM unlabeled protein, Light grey circles indicate 50nM labeled protein in 10µM unlabeled protein, and Deep grey circles indicate 50nM labeled protein in 20µM unlabeled protein. All measurements were take n=3. The data are shown mean ± SD.

The variations of r_Hm_ with the concentrations of Dextran 70 under these four different protein concentrations (C-I to C-IV) are shown in Figure 3c. Figure S5a and Figure S5b show the same for Dextran 40 and Dextran 20. We calculated the extent of change in r_Hm_ for each condition as Δr_Hm_, which is the difference between r_Hmnc_ and r_Hmc_, where r_Hmnc_ is the value of r_Hm_ in the absence of any crowding agent and r_Hmc_ is the value of r_Hm_ at any crowder concentration (see Figure 3d). We would refer to the concentration of the crowder where r_H_ is the minimum as C_comp_ (please refer to Figure 3d). The variation of Δr_Hm_ with the concentrations of Dextran 70 under conditions between C-I and C-IV are shown in Figure 3e. The corresponding data for Dextran 40 and Dextran 20 are shown in Figures S5c-d.

The data for all these Dextrans (Figure 3c,e; Figure S5a,b; and Figure S5c,d for Dextran 70, 40 and 20) clearly showed a bi-phasic behaviour between a native-like state (N), compact state (C) and an extended state (U). The first transition (compaction or the transition from N to C) occurs at a relatively lower Dextran concentration. This was followed by an increase in r_Hm_ (the formation of U from C), which occurred at a relatively higher concentration of Dextran.

We found out that both the extent of compaction decreased when we increased the protein concentrations (the representative Figure 3f). Since Δr_Hm_ is most prominent in the absence of any unlabeled protein, we used this experimental condition to investigate the effect of compaction in more detail. We found that at this condition (without unlabelled protein) the extent of compaction (Δr_Hm_)_Ccomp_ increased with the increase in the molecular weight of Dextran; i.e., the extent of the compaction followed the following trend: Dextran 70> Dextran 40> Dextran 20 (Figure 3f, inset). To check it further, we also used Dextran 6, which showed no compaction and expansion (Figure 3g). To determine the extent of compaction and/or expansion varied for Ficoll-70, we used FCS experiments with labeled Sup35NM in the absence and presence of different concentrations of Ficoll-70. Interestingly, although we observed the formation of the compact state in the presence of Ficoll 70, this crowder did not yield the formation of the extended state (Figure S5g). The variable effect of Dextran and Ficoll on Sup35NM, which we observed experimentally, prompted us to computationally model and test the effect of these two at single-molecule level. In this attempt, a set of initial models of Sup35NM were computationally built from its sequence information using the *de-novo* modeling approach of the iTASSER program^1^. In order to afford long time-scale simulation, two top-scoring predicted models of Sup35NM were converted to a coarse-grained resolution by applying the Martini 3 model and multiple Molecular Dynamics (MD) simulations were performed to populate a cumulative ensemble of 30 *μ*s. The distribution of the radius of gyration (R_g_) of the protein (Figure S6a) ensemble covers a broad range, from 2 to 6 nm, indicating that Sup35NM monomer populates a diverse conformational ensemble consisting of compact to extended conformations. Further, using the *k*-means clustering approach, we clustered the broad protein ensemble into three states, S1 to S3, (Figure 4a,b) that is consistent with the experimentally observed native-like (N), compact (C) and extended states (U), respectively. A representative conformation from each cluster/state was then used to study the effect of a crowded environment. To mimic the extended (Dextran) and spherical (Ficoll) shape of the crowding agents used in experiments, we modelled an extended rod-like polymer and a spherical crowder with C60 fullerene-like topology. The crowders were modelled to be inert, thereby exerting only the excluded-volume effect. We studied the effect of these crowding agents at two concentrations, i.e. 10% v/v and 30% v/v of the simulation box, on each of the three conformational states of Sup35NM. We performed three simulations (10 *μ*s each run) for each conformation in the presence of the two types of crowders and from the cumulative simulation data for each case, we estimated the degree of change in the extent of compaction induced by the excluded volume effect of the crowders. The degree of change of R_g_ of the protein with respect to its initial R_g_ is shown in Figure S6b. In the presence of spherical crowders, the protein undergoes compaction and the extent of compaction is proportional to the R_g_ of the initial structures. We found that the most extended initial structure undergoes maximum compaction. Furthermore, the percentage compaction increases with increasing crowder concentration. Sup35NM attains a mean R_g_ value of ∼2.0 nm and ∼1.7 nm in the presence of 10 % v/v and 30 % v/v spherical crowder concentrations, respectively. On the other hand, the presence of extended crowders in the solution has an interesting effect on the protein conformation. While N and U initial states of Sup35NM (corresponding to S2 and S3, respectively) undergo compaction in the presence of extended crowders, the most compact state (C) unfolds and expands. We note that the degree of compaction of the protein is significantly lower in case of extended crowders when compared to spherical crowders (see Figure S6b), which supported our experimental results.

**Figure 4:**
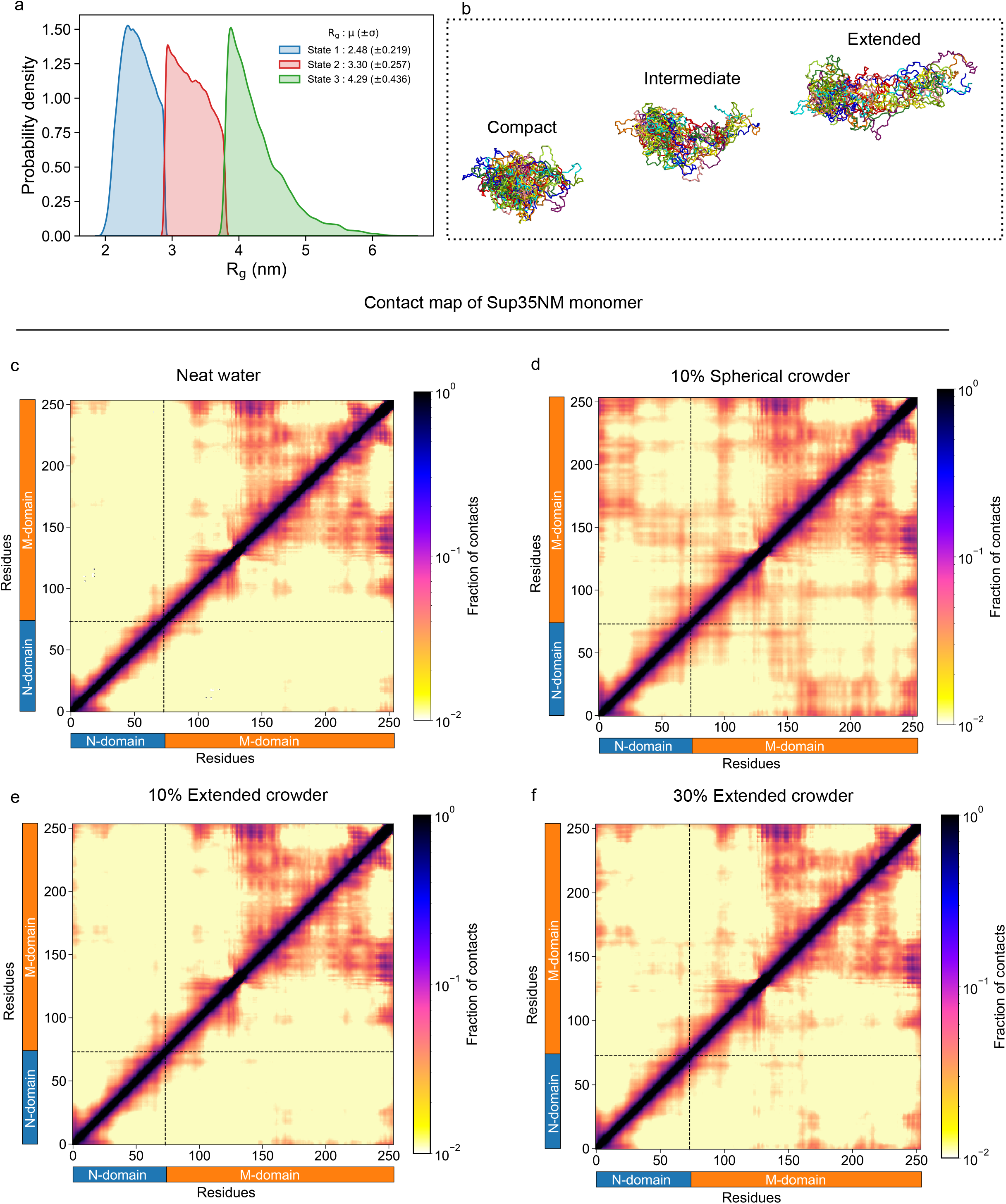
Coarse-grained simulations of Sup35NM monomer: Probability distribution of R_g_ of the ensemble (a) k-means clustering of Sup35NM monomer ensemble based on R_g_ into three states. The mean and standard deviation of R_g_ of the three states is provided in the legend (b) Representative snapshots obtained from coarse-grained simulations of overlay of multiple conformations for compact, intermediate and extended conformations. (c) Average intraprotein residue-wise contact probability maps of Sup35NM monomer in neat water (d) Average intraprotein residue-wise contact probability maps of Sup35NM monomer in presence of 10 % v/v spherical crowder, (e) Average intraprotein residue-wise contact probability maps of Sup35NM monomer in 10 % v/v extended crowder, (f) Average intraprotein residue-wise contact probability maps of Sup35NM monomer in 30 % v/v extended crowders. Axes denote the residue numbers. The colour scale for the contact probability is shown at the extreme right of each panel of maps. The colour bar along the axes of the plots represents the domains in the protein.

Next, we compared the structural properties of the conformational ensemble of Sup35NM in the absence and presence of crowded environments. We estimated the average inter-residue contact probabilities of the ensemble in neat water (in the absence of any crowder) and in presence of the crowders (Figure 4c-f, S6c). The contact map of the neat water ensemble (Figure 4c) indicates that the conformations populated are characterized by intra-domain interactions within the M-domain. These contacts are present between residues ranging from ∼80 to 150 (consisting of uncharged polar amino acids, namely, glutamine and asparagine) and the C-terminal part of M-domain comprising acidic amino acids (glutamic acid and aspartic acid). These intra-domain interactions within the M-domain are also prevalent in the presence of crowders both in the presence of 10% and 30% crowder concentrations (Figure 4d-f, S6c).

The presence of spherical crowders lead to significant protein compaction, and these compact conformations are characterized by both intra- and inter-domain interactions (Figure 4d, S6c). Along with the intra-domain interactions within the M-domain, long-range interactions are predominantly formed between the N-domain and the residues of the M-domain (residues # 150 to 253). At higher concentration (30% v/v) of the spherical crowder, the inter-domain interactions are significantly enhanced (Figure S6c). However, in the presence of extended crowders (Figure 4 e,f), the long-range interactions between the N (∼residues 1 to 20) and M domains (∼residues 150 to 253) are present to a much lesser extent than that of spherical crowders. This can be attributed to the dual effect i.e. protein expansion and compaction, promoted by volume exclusion of extended crowders in the solution. These observations highlight the propensity of inter-domain long-range interactions in the conformations favoured under crowded environments.

### Compaction, expansion, and Phase Transition

Previous SANS data and coarse-grained simulations have shown the presence of crowder-induced compact and expanded conformations in FlgM, an intrinsically disordered protein involved in transcriptional regulation in *Salmonella typhimurium* ^29^. We found that the extent of compaction (or expansion) depends on the molecular weight of Dextran. We also found that in the presence of Dextran 6, there was no change in hydrodynamic radius of SUP35 presumably reflects its comparable hydrodynamic radius with the protein (the protein SUP35, R_H, SUP35_ ∼ 18.4Å with dextran 6, R_H, Dextran-6_ ∼18Å)^26^. The initial reduction of protein hydrodynamic radius up to a specific protein, crowder concentration may be due to crowder induced steric repulsion and localization of protein conformers in interstitial void volume. The degree of compaction was higher in dextran 70 and lower in dextran 20 may be attributed to the pressure exerted by the crowder molecule on a protein molecule owing to their molecular weight resulting in more intra-domain interaction of IDP^26^. Another interesting observation is that with Dextran concentration less than C_comp_ (where the protein conformation is predominantly compact) there is no phase separation. In contrast, beyond C_comp_, where the protein samples adopted the extended U state, the initiation of phase separation becomes apparent. It may be noted that depletion interactions/force has been recently suggested to be one of the key factors that lead to “phase separation” of protein molecules. It is possible that the conformational expansion observed here may create a larger depletion layer around the protein molecule. The depletion layer thickness is directly proportional to the hydrodynamic radius of the polymer crowder^30^. Beyond C_comp_, depletion attraction creates an associative interaction that favours LLPS^28^. We also found that an increase in protein concentration shifts protein phase separation towards a lower crowder concentration. At high protein concentrations (20 *μ*M or beyond), more protein molecules are available per unit volume to overlap with each other. A considerable amount of overlap between depletion layers is thus possible even at low crowder concentrations.

Previous studies on expansion and compaction attribute the phenomena to soft interactions between a protein and crowding agents in addition to the effect of crowder shape^31,32^. The inertness of the crowder, modelled in the present simulations, was manifested as hard-core steric repulsions between the protein and crowder molecules. Using this strategy, we were able to recapitulate the experimental observation of both compact and extended states of the proteins in the crowded environment. Here, the key determinant in influencing the protein conformational states is the shape of the inert crowder. The spherical crowders exert their effect by stabilizing the compact state of the protein in the interstitial void. On the other hand, the rod-like extended crowders stabilize the protein state compatible with the corresponding void volume. In the process of attaining the stable state, the protein may undergo expansion by winding through the interstitial crevices in highly crowded solution, or in order to fit the void in a relatively less crowded solution, may experience compaction^29^.

To determine the molecular drivers that govern aggregation and phase separation of Sup35NM and how this process is modulated in a crowded environment, we performed coarse-grained simulations of multiple chains of Sup35NM in neat water and in the presence of crowders. In these simulations, polydispersity of the initial protein conformations was maintained using the ratio of the three states S1 to S3 (compact, intermediate and extended conformations) which was identified in the parent ensemble (see Figure 4a). To detect aggregation events, we estimated the number of monomers and the cluster/aggregate sizes that are formed in the simulations. The distribution of the number of monomers in the solution with their mean and standard deviations is depicted in Figure 5a. In the presence of spherical crowder, the number of monomers was found maximum. The cluster size distribution (Figure 5b) was observed to be the broadest (ranging from dimer to 11-mers) for neat water conditions when compared to that observed in the presence of crowders. The aggregate sizes varied between dimer and 9-mers for the extended and between dimer and 10-mers for the spherical crowders, respectively. These observations that Sup35NM would aggregate more in the presence of spherical crowders when compared to the extended crowders, which is consistent with the experimental observations, as phase separation was completely absence in the presence of Ficoll.

**Figure 5:**
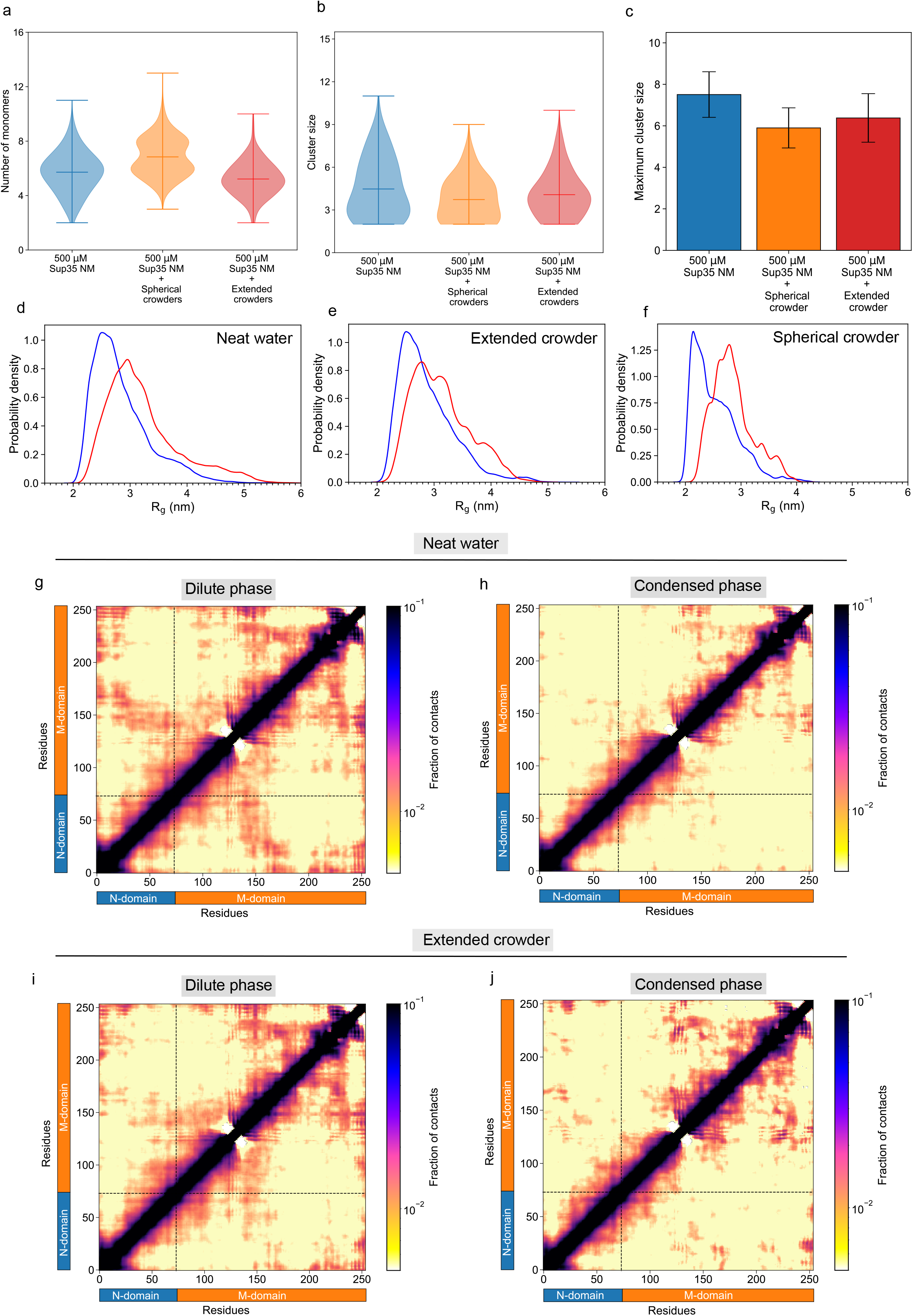
Aggregation simulation of Sup35NM in different crowding agents: (a) Distribution of the number of monomers in aggregation simulations in the presence and absence of crowders, (b) Distribution of the cluster sizes and (c) Bar plot of the maximum size of the aggregates. All measurements were taken in triplicate, n=3. The data are shown mean ± SD. Probability distributions of R_g_ of the protein in the monomer phase and aggregates in the (d) neat water and in presence of, (e) extended crowders and (f) spherical crowders. Average intraprotein residue-wise contact probability maps of Sup35NM in the (g) dilute and (h) dense (aggregate) phase in (g) neat water. Average intraprotein residue-wise contact probability maps of Sup35NM in the (i) dilute and (j) dense (aggregate) phase in extended crowders. Axes denote the residue numbers. The colour scale for the contact probability is shown at the extreme right of each plot of maps. The colour bar along the axes of the plots represents the domains in the protein.

Emergent evidence suggests that upon phase separation, IDPs adopt more elongated or extended conformations^2,3^. To verify if the aggregated protein chains are more extended than free monomers, we quantified the R_g_ of all protein chains and compared their probability distributions for the dilute and dense phase in each system, as illustrated in Figure 5d,f. Here, we considered molecules with molecular weights of hexamers and beyond as the components of the dense phase. For both in the absence (neat water) and presence of the crowder, we found that the probability distributions for the dense phase were broader and shifted towards higher values. The results implicate that the protein chains in aggregates adopt more extended conformations when compared to chains in the dilute phase. Among the three ensembles, the monomers and aggregates in neat water ensemble show the highest difference in R_g_ distribution. They also have the broadest distribution and non-overlapping peaks with dense phase R_g_ ranging from 2 to 6 nm while neat water R_g_ ranging from 2 to 5 nm. Among the two types of crowders, the spread of the R_g_ distributions of the protein in presence of spherical crowders is from 2 to 4 nm, with the distinct dominant peaks at ∼2.1 nm and ∼2.9 nm for chains in net water and dense phase, respectively. Furthermore, in the presence of extended crowders, the distributions spread across 2 and 5 nm. These observations are consistent with previous experimental and computational studies that report favourability for extended states in the dense phase as a characteristic hallmark of liquid-liquid phase separation^33^.

In the light of these observations, we delved into a residue-level investigation of the intra- and inter-domain contacts, which may be responsible for the stabilization of the dense phase formed in the presence and absence of the crowders. Figure 5g-h, S7 depicts the average intra-protein contact maps in the three ensembles (neat water, spherical crowders and extended crowders) in the dilute and dense phases. In general, the contact maps illustrate a diverse network of intra- and inter-domain interactions characterizing these states^2^. The protein chains that are in dilute phase in the presence of spherical crowders (Figure S7a) are marked by the densest network of intra-domain interactions. Importantly, long-range interactions are formed between the N-domain that is enriched in polar residues asparagine, glutamine and tyrosine and the M-domain regions residues (140 to 160 and 200 to 250). There are mostly charged residues, namely, lysine, aspartic acid and glutamic acid. The presence of long-range contacts indicates that these monomeric conformations are highly compact in nature. This is consistent with our experimental observation of the decrease in hydrodynamic radius with spherical Ficoll crowders in the solution (see Figure S5g) as well as our monomer simulations with spherical crowders (see Figure 5g-h, S7). The contact maps of protein chains in the dilute phase in neat water as well in presence of extended crowders (Figure S6g,i) also show intra-domain interactions in the M-domain and long-range interactions between the two termini. However, the extent of these interactions is reduced when extended crowders are present. In the dense phase, the extended nature of the protein chains compared to the chains in dilute phase is clearly reflected in the intra-protein contact maps (Figure 5h,i, S7b). The least protein expansion is observed in the ensemble with spherical crowders and therefore the corresponding average contact map (Figure S7b) shows intra-domain long-range interactions. Relatively less extent of long-range interactions is also observed in the protein chains of the dense phase in neat water (Figure 5h). Importantly, in the presence of extended crowders, the protein conformations that form the dense phase are most extended in nature (Figure 5j).

To elucidate the inter-chain interactions that stabilize the dense phase, we analysed the residue-residue interactions between the protein chains of the neat water and crowder ensembles (Figure S8). The average contact maps are evidently distinct for the three ensembles. In the dense phase formed in neat water (Figure S8a), the polar, uncharged residues of the N-domain form interactions with the residues across the entire M-domain consisting of polar residues. The contacts are particularly prominent between polar, uncharged residues of N-domain with charged residues 200 to 250 of the M-domain. In the aggregates formed with extended crowders in solution (Figure S8b), long-range interactions between the N-domain with charged residues of the C-terminal (residues 200 to 250) are dominant. Moreover, polar, uncharged residues of the M-domain also interact with the charged residues 200 to 250. Lastly, In the aggregates formed when spherical crowders are present, the predominant interactions are between the N-domain with residues in the range ∼90 to 150 in the M-domain (Figure S8c). These observations described above from the aggregation simulations shed light on the modulatory effect that crowders have on the aggregation propensity and nature of the resultant aggregates of Sup35NM (Figure S8d-f). Furthermore, the residue-level analyses provide insights into the nature of the interactions that govern the oligomers formed in the presence and absence of crowding agents in solution.

## Conclusions

Although molecular crowding has been suggested to play crucial roles in protein phase separation, systematic studies of the effect of the shape and size of the crowder molecules have been limited. In this study, using single-molecule fluorescence correlation spectroscopy along with computational methods we showed that an intrinsically disordered protein, like Sup35NM, can adopt multiple conformational states depending upon the size, shape and molecular weight of crowder molecule. We have shown here that up to a specific crowder concentration steric repulsion and excluded volume effect stabilizes the compacted population of IDPs but as we kept on increasing the chain length, branching and concentration of crowder molecule it affected the microviscosity and translational diffusion of IDP and depletion interaction started to prevail which helps in the associative attraction of protein molecule leading to formation of distinct biomolecular condensate. Using FCS, we also tried to correlate the phase separation behavior of IDP with their population distribution of different conformers in cellular-like crowded environment. We showed that the transition between N↔C↔U of IDP is highly dependent upon the physical properties of the crowder molecule. Altogether in our study, we tried to find a correlation between spatial IDP conformations to their propensity of biomolecular condensate formation in cellular-like crowded environments using synthetic crowders (Figure 6). The considerable dependence of crowding effects on IDP can be explained by depletion force and polymer theory. An interesting finding of our study is how associative kinetics balance the viscosity induced resistance of diffusion and tried to stabilize the protein in liquid droplets. Our study can be helpful for further exploration of finding out a universal model system which would be able to sufficiently diagnose the different conformational landscapes of IDPs in the presence of wide range crowder molecules mimicking the cellular environment which will help in solving the functioning and disease correlation of IDP in our cells.

**Figure 6:**
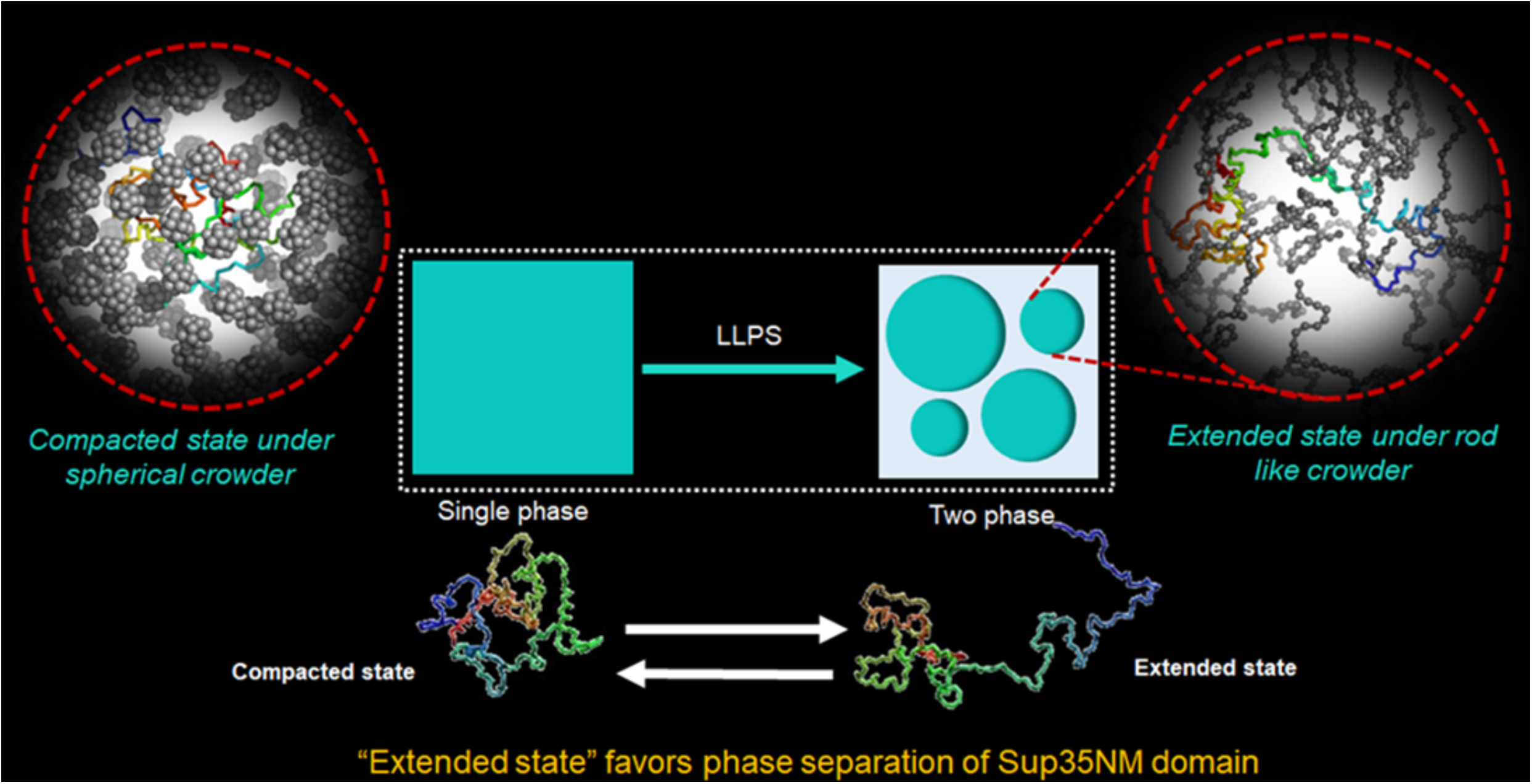
Schematic representation of conformationally guided phase separation of Sup35NM in a crowded milieu: Sup35 can exhibit both compacted and extended conformers in the presence of extended (rod-like) crowder (like Dextran). The extended/elongated conformer favours droplet formation. The droplet formation is not favoured by the compact conformers (predominant in the presence of spherical crowder, like Ficoll) of Sup35NM.

## Acknowledgements

We thank Sounak Bhattacharya for technical support in confocal microscopy and Central Instrumentation Facility, Indian Institute of Chemical Biology for the provision of infrastructure. The work done at the Indian Institute of Chemical Biology was supported by the fund MLP 139. JM acknowledge the support of the Department of Atomic Energy, Government of India, under Project Identification No. RTI 4007 and Core Research grants provided by the Department of Science and Technology (DST) of India (CRG/2023/001426).

## Author Contributions

SR and NM carried out the experiments and analyzed the data. SM performed and analyzed the simulation data. KC and JM supervised the study. SR, SM and NM wrote the initial draft of the manuscript. KC and JM prepared the final version. All authors approved the final version of the manuscript.

