## Supporting Information for "Crowder induced conformational fluctuations modulate the phase separation of yeast SUP35 NM domain"

### Materials and Methods

#### Purification of protein

Recombinant SUP35NM protein was expressed and purified following the procedure of *Glover et al*. with slight modification[1]. The plasmid for SUP35NM expression in E.coli cells (BL21 DE3 strain) pJCSUP35NM (1-253) was obtained from ADDGENE containing plasmid number #1089. The cells were induced using 1M IPTG for 3.5-4 h for over expression of the protein. Harvesting of the bacterial cells was performed at 6000 rpm at 4°C for 15 min. Purification of SUP35NM was performed under denatured conditions at ambient temperature. The cells were lysed by gentle homogenization for 30 min using pre-chilled lysis buffer (20mM Tris-HCl, 8M urea, pH 8.0). After that the insoluble cell debris was pelleted down by centrifugation at 10,000 rpm for 30 min at 4°C and the supernatant was separated. The soluble protein portion was mixed with sufficient amount of Ni-NTA and kept at 4°C for 4h with gentle shaking for proper binding. The slurry of Ni-NTA bounded protein was poured in a column and washed using ~5 bed volume of wash buffer (50 mM imidazole, 20mM Tris-HCl, pH 8.0) and eluted with ~2 bed volume of elute buffer ( 20 mM Tris HCl, 500 mM imidazole at pH 8.0). The eluted portion was collected and dialyzed in 20mM phosphate buffer solution (PBS) of pH 7.4 for overnight at 4°C using snake skin dialysis membrane of molecular cut-off 10KD. The final presence of protein molecule was confirmed by SDS-PAGE gel electrophoresis and MALDI. Final protein concentration was evaluated using molar extinction co-efficient of 29,000 $M^{-1}$ $cm^{-1}$ at 280 nm wavelength for the monomeric protein using UV-VIS spectroscopy.

#### Fluorescent labelling of protein

Purified Sup35 protein was labeled using Rhodamine Green™-X, Succinimidyl Ester, Hydrochloride mixed isomers (Invitrogen, MA, USA) following previously established protocol. The fluorescence dye was dissolved in DMSO and mixed with a 2mg/mL solution of protein under continuous stirring. The molar ratio between the protein and dye was 1:10. This reaction mixture was incubated at 4 °C for 5 hours with a vortexing post every 30 minutes. The labeling reaction was then quenched by adding excess β-mercaptoethanol. Excess free dye from the reaction mixture was removed by extensive dialysis in Na-phosphate (pH 7.4) buffer using SnakeSkin (Thermo Fisher Scientific, MA, USA) Dialysis Tubing (10 kDa MWCO) followed by column chromatography using a Sephadex G20 column (Bio-rad Laboratories, CA, USA) pre-equilibrated with 20 mM Na-phosphate buffer (pH 7.4)[2].

#### In-vitro phase separation assay

The lyophilized protein sample was dissolved in NaP (pH= 7.4) buffer and concentration was measured using a UV-VIS spectrophotometer. Phase separation reaction mixtures were prepared by mixing unlabelled protein at different concentrations along with 1% labeled protein, 100 mM NaCl and the required percentage of Dextran/Ficol variants according to the experimental conditions. For incubation periods and time points, the Eppendorf tubes were kept in a moist chamber at 37°C with shaking at 180 rpm. The phase separation was confirmed by turbidity measurement (Absorbance at 600 nm) in UV-VIS spectrophotometer and corresponding images were taken in a confocal or fluorescence microscope in DIC or fluorescence mode[3].

**Fluorescence Correlation Spectroscopy (FCS)**

Fluorescence Correlation Spectroscopy (FCS) measures the fluctuations of fluorescence intensity in the confocal volume and yields the diffusion times of the fluorescent species. The FCS measurements were carried out using an ISS Alba FFS/FLIM confocal system (Champaign, IL, USA), coupled to a Nikon Ti2U microscope equipped with the Nikon CFI Plan Apo 60X / 1.2NA water immersion objective. The 488-nm picosecond pulsed diode laser was used for the excitation of the FCS measurements. The fluorescence emission was recorded using a pair of SPAD (Single Photon Avalanche Detector) detectors with the 50/50 beam splitter and the 530/43-nm band-pass filter. By utilizing two detectors, we were able to compute cross-correlation functions for individual colors, thereby eliminating any artifacts caused by the after-pulsing of the detector. The FCS correlation curves were fit to the 3D Gaussian 1-component diffusion model, where the beam waists in the radial and axial dimensions were calibrated using a standard fluorescence dye (Rhodamine 6G, R6G) of known diffusion coefficient (2.8 × 10−6 cm2/sec). The diffusion coefficient of different mutants is measured by measuring the diffusion time ($\tau_D$) and using the relation:

$$\tau_D = {r_0}^2/4D$$

Where $r_0$ is the lateral radius of the hypothetical ellipsoid observed volume. The hydrodynamic radii of each species were computed from their respective diffusion coefficients (D). For comparison between hydrodynamic radii of the wild-type protein and its mutants, Alexa Fluor 488 labeled monomeric protein (50μM and 100 nm unlabeled protein in the presence of 10 mM TCEP in 20 mM Na-phosphate buffer (pH 7.4) was used. The recorded FCS data were normalized with respect to free dye following a previously published method 24 In the 3D Gaussian diffusion model involving a single type of diffusing molecules (the 3D Gaussian 1-component model eliminating the contributions of the triplet state), the correlation function G (τ) can be described by the following equation:

$$G(\tau) = \frac{1}{N}.(1+\frac{\tau}{\tau_D})^{-1}.(1+\frac{\tau}{S^2\tau_D})^{-1/2}\left[1+K_{exp}\left(-\frac{\tau}{\tau_S}\right)\right] \ldots\ldots\ldots\ldots\ldots(13)$$

Where N is the number of molecules, $\tau_D$ is the diffusion time and S is the ratio between axial and lateral radius, Kexp is the chemical relaxation constant at equilibrium condition and $\tau_S$ is the relaxation time. The value of D calculated by fitting the correlation function is related to the diffusion coefficient (D) of a molecule by the equation below:

$$\tau\mathrm{D} = \frac{\omega^2}{4D}$$

Where ω is the size of the confocal volume. From here, the value of the hydrodynamic radius (rH) of the fluorophore-labeled molecule can be computed from D using the Stokes-Einstein formula:

$$D = K.T/6\pi\eta r_H$$

Where η is the viscosity, T is the absolute temperature and k is the Boltzmann constant[4,5].

### Thioflavin-T (ThT) binding assay

For aggregation monitoring, 500 µL of 20 µM of protein in NaP was incubated in the absence and in the presence of different concentrations of Dextran-70. 1 mM sodium azide was added to prevent any bacterial contamination in the reaction mixture. This solution was kept under 180 rpm and 37°C shaking conditions during the aggregation kinetics. The aliquots were withdrawn at different time points and diluted to a volume of 500 µL in NaP buffer to keep a working concentration of 2 µM for Sup35 protein. The protein: ThT was kept at 1:10 for fibrillation pathway determination. All measurements were recorded in a PTI Quantamaster spectrofluorimeter (Horiba, Japan) using a quartz cuvette of 1cm path length. The steady-state fluorescence of ThT at 485 nm was measured after proper mixing, using excitation at 440 nm. The average of 3 measurements was taken for analysis. The excitation and emission slit widths were 5 nm and the integration time was 0.1 s. For samples that gave sigmoidal kinetic profiles, the Boltzmann equation (Eqn. 1) was used for fitting.

$$y = y_0 + \frac{A}{1+\exp\,(k'(t-t_{0.5})} \quad \ldots\ldots\ldots\ldots\ldots (1)$$

$y_0$ = baseline signal, A= total increase in fluorescence, k' = growth rate constant, $t_{0.5}$ = midpoint of the transition from which the lag time ($t_{lag}$) can be calculated by using the following Eqn. 2

$$t_{lag} = t_{0.5} - \frac{1}{2K} \ldots\ldots\ldots\ldots (2)$$

$t_{lag}$ = time point where the amplitude of the transition reaches 10% of the whole transition[6,7].

### Nuclear Magnetic Resonance (NMR) experiment

$^1$H NMR Spectra of SUP35 and SUP35 + Dextran 70 were recorded on Bruker Avance 600MHz spectrometer. The experiment was performed in D2O taking tetramethylsilane (TMS) as standard molecule. 32 repeated scans were taken and the data were reported in chemical shifts (ppm) relative to TMS.

### Atomic force microscopy (AFM)

Aggregated samples of Sup35 were aliquoted after prolonged incubation at 37 °C following 20 times dilution with MilliQ water. On freshly cleaved mica, a 5 µL diluted sample was drop cast. After rinsing the aggregates with MilliQ water, they were dried using a stream of nitrogen. Using a Bioscope Catalyst AFM

(Bruker Corporation, MA, USA) with silicon probes, images were acquired at room temperature. To analyse the morphological features of aggregates, a standard tapping mode was utilized. The nominal spring constant of the cantilever was kept at 20–80 N/m. The spring constant was calibrated by a thermal tuning method. A standard scan rate of 0.5 Hz with 512 samples per line 6 was used for imaging the samples. Following a single third-order flattening of height images with a low pass filter, section analysis was performed to determine the dimensions of aggregates[2].

**Laser scanning confocal microscopy (LSCM)**

Droplet formation by Sup35NM domain was confirmed by laser scanning confocal microscopy along with the turbidity measurement. The unlabeled protein was doped with 15nM label protein to visualize in the microscope. Then the samples were prepared using different dextran concentrations ranging from 0mg/ml to 100mg/ml. The concentration range was same for all dextran variants. The samples were then incubated for ~30 mins and drop-casted in a glass slide (Blue Star, India) using a pipette and it was mounted by 22 mm coverslip (Blue Star, India). The samples were sealed using locally available nail paint. Then it was imaged using a Leica SP8 (Leica, Germany) laser scanning confocal microscope using a 488 nm excitation laser and suitable detectors. For the temporal maturation study with dextran-70 the samples were taken out at different time points and imaged in a similar manner using Leica SP8 (Leica, Germany) laser scanning confocal microscope[7].

**Computational study of Sup35NM**

**Coarse-grained simulations of Sup35NM monomer in neat water**

The NM domain of the yeast prion protein Sup35 (Sup35NM) that regulates the prion behaviour of the protein was used in this study. Since the structure of the Sup35NM is unknown, we used an ab-initio protein structure modelling and refinement approach known as I-TASSER. We chose 2 models out of the 5 top-scoring models predicted by I-TASSER for the simulation study. Using a similar approach, Sup35NM model structure was used in a recent study of the role of membrane composition. These three model structures were converted to coarse-grained resolution of Martini 3 3 model for biomolecular simulations using Martinize24. The protein conformation was solvated in CG Martini water and the system was made electroneutral. The CG simulations were performed using the GROMACS simulation package5. The resulting simulation system was energy minimized for 10000 steps using the steepest descent algorithm. This was followed by equilibration in the NVT ensemble at a time step of 10 ps for a timescale of 5 ns. Another round of equilibration in the NPT ensemble was performed at a time step of 20 ps for 10 ns timescale. The v-rescale thermostat6 was used with a coupling time of 1 ps to maintain the temperature at 298.15 K. Constant pressure was maintained at 1 bar using the Berendsen barostat7 with a relaxation time of 12 ps. Nonbonded interactions were cut-off at 1.1 nm and Coulombic interactions were calculated using the Reaction Field method8. Production runs were performed on the CG system using a time step of 20 ps for 10 μs timescale[8].

The ensembles of conformations obtained from the multiple simulations were subjected to k-means clustering using radius of gyration (Rg) as the collective variable. The conformations were segregated into 3 clusters corresponding to the number of conformer populations obtained in experiments. The Rg corresponding to these clusters are 2.48 (± 0.219) nm, 3.30 (± 0.257) nm and 4.29 (± 0.436) nm, respectively.

**Simulations of Sup35NM monomer in the presence of model crowders**

A representative conformation was obtained from each of the 3 clusters and to study the effect of a crowded environment. Two types of model crowders varying in shape were considered: spherical and extended crowders. The spherical crowder was modelled using the topology of C60 fullerene. The CG representation and parameters of the fullerene crowders was obtained from the GitHub repository (https://github.com/JMLab-tifrh/fullerene-Martini3_repulsive)[9]. The polymer crowded was modeled using the Martini model of polyethylene polymer. The polyply program was used to build the polymer model. Since polyethylene is a hydrophobic polymer, neat water is not a good solvent and the polymer chain collapses. To create an extended polymer chain, we tuned the epsilon (ε) nonbonded parameters of C1 (Martini apolar bead type for butane) with water to make the interaction favourable. A ε value of 5.5 caused the polymer to favourably interact with water and remain extended yet flexible. We also made the polymer inert by setting a negative σ value of the nonbonded interactions (repulsive interaction) of C1 bead with all other beads types.

The model crowders were rendered inert with respect to the protein atoms and other crowder molecules by introducing repulsive interactions between these components. However, the interactions with the solvent and ions were maintained as the full potential of attractive and repulsive interactions. The crowding simulation setup (consisting of the protein, crowders, solvent and ions) was prepared at 2 crowder concentrations: 10 % v/v and 30 % v/v of the simulation box. For each of the 3 protein conformations corresponding to the three clusters, three replicates of the CG simulations were performed for a timescale of 10 μs[10,11].

**Aggregation simulations of Sup35NM**

Multi-chain simulations were performed to study the effect of crowders of different shapes (spherical and elongated) on the aggregation propensity of Sup35NM. In these simulations, the polydispersity of the initial protein conformations was ensured by maintaining the same population of the three states C1 (53.26 %), C2 (32.68%) and C3 (14.05 %) as present in the neat-water ensemble that was subjected to clustering based on Rg. Considering the limitations of computational time, 30 protein chains amounting to 500 $\mu$M concentration (higher than the experimental concentration) was used for the aggregation simulations. Two types of crowding simulations were prepared, corresponding to spherical and extended crowders, respectively. Similar to monomer simulations, a crowder concentration of 10 % v/v was added. A control simulation of multiple protein chains in neat water was also prepared. These systems were then solvated with CG Martini water and 100 mM NaCl (as in experiments) was added. The simulation details are the same as described for monomer simulations above. In each case (i.e. neat water, spherical crowders and extended crowders), two simulations were performed for a timescale of 2 $\mu$s. For analyses, the last 1 $\mu$s of the simulation trajectories was considered[12].

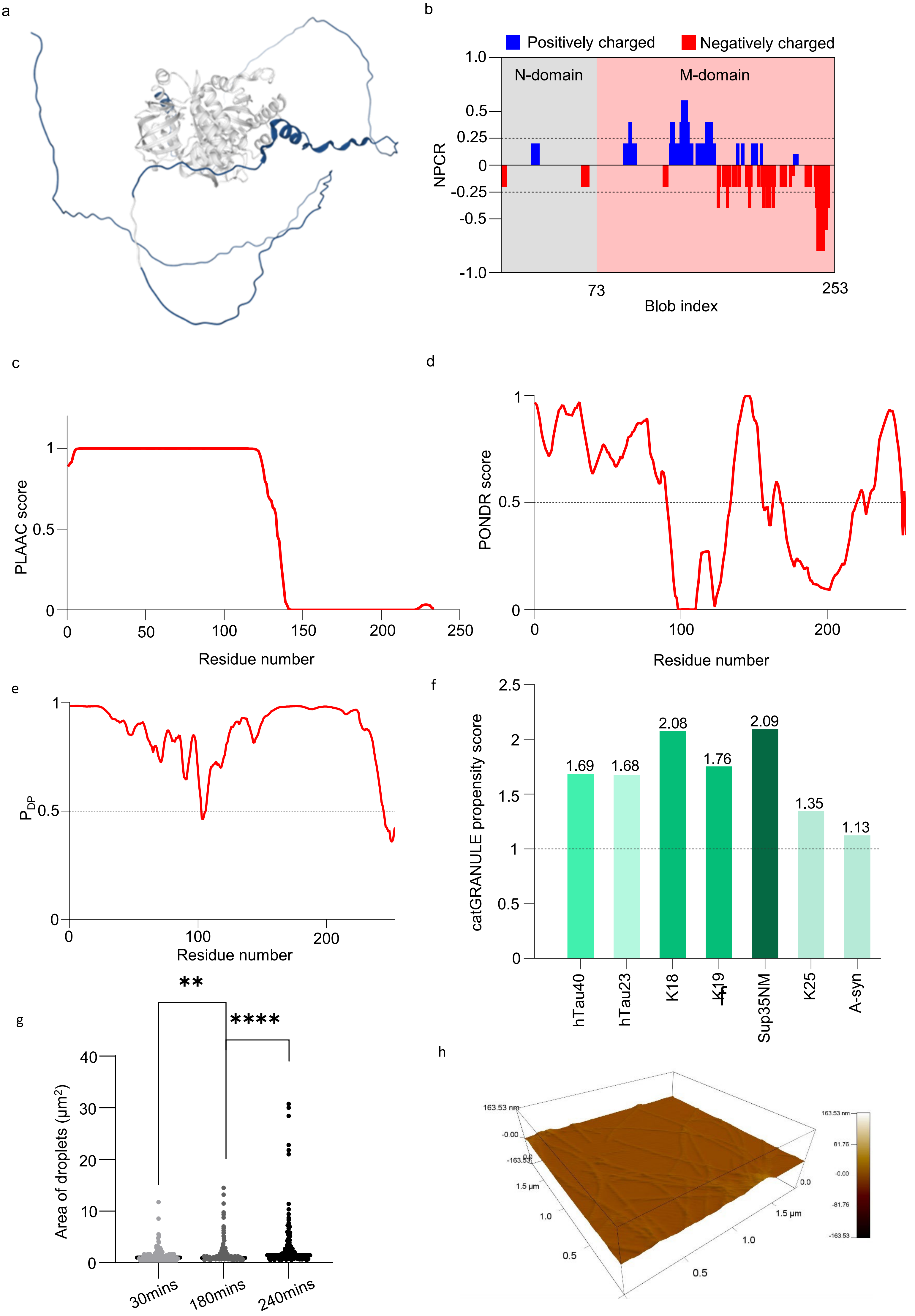
a
b
Positively charged
Negatively charged
N-domain
M-domain
NPCR
1.0
0.5
0.25
0
-0.25
-0.5
-1.0
73
253
Blob index
c
PLAAC score
Residue number
0
50
100
150
200
250
d
PONDR score
Residue number
e
$P_{DP}$
Residue number
f
catGRANULE propensity score
1.69
1.68
2.08
1.76
2.09
1.35
1.13
hTau40
hTau23
K18
K19
Sup35NM
K25
A-syn
g
**
****
Area of droplets (μm²)
40
30
20
10
0
30mins
180mins
240mins
h
163.53 nm
-0.00
-163.53
1.5 μm
1.0
0.5
0.0
81.76
-81.76

**Figure S1**: (a) Model structure of full-length yeast prion Sup35 from AlphaFold. (b) Net charge per residue (NPCR) plot of Sup35NM domain calculated from CIDER tool. The blue colour indicates the positively charged amino acids and the red colour indicates the negatively charged amino acids. The M-domain contains more charged residues compared with the N-domain. (c) Prion-like amino acid propensity was measured from the linear sequence using the PLAAC tool. The aggregation-prone, prion-forming N-domain and portion of the M-domain have prion-like amino acid composition. (d) Disorder prediction from Predictor of Natural Disordered Regions (PONDR) indicates the disorder region in the sequence (Above 0.5 threshold indicates the disorder region). The regions of amino acids 1-91, 134-155, 164-168, and 220-250 were found to be disordered. (e) The probability of phase separation from the linear sequence of Supo35 using FuzPred indicates the entire NM region of the protein has a high propensity for phase separation. (f) Comparison of catGRANULE scores of different phase separating proteins, full-length tau (hTau40), repeat domains of Tau (K18 and k19) along with Sup35NM. (g) Droplet Size distribution of temporal maturation indicates an increase in the average size of the droplet with time. Three independent measurements were taken. Statistical significance was measured using paired t-test; **$P < 0.01$; ****$P < 0.0001$. (h) 3-D plot for measuring height distribution of the Sup35NM fibrils as represented in Figure 1h, inset.

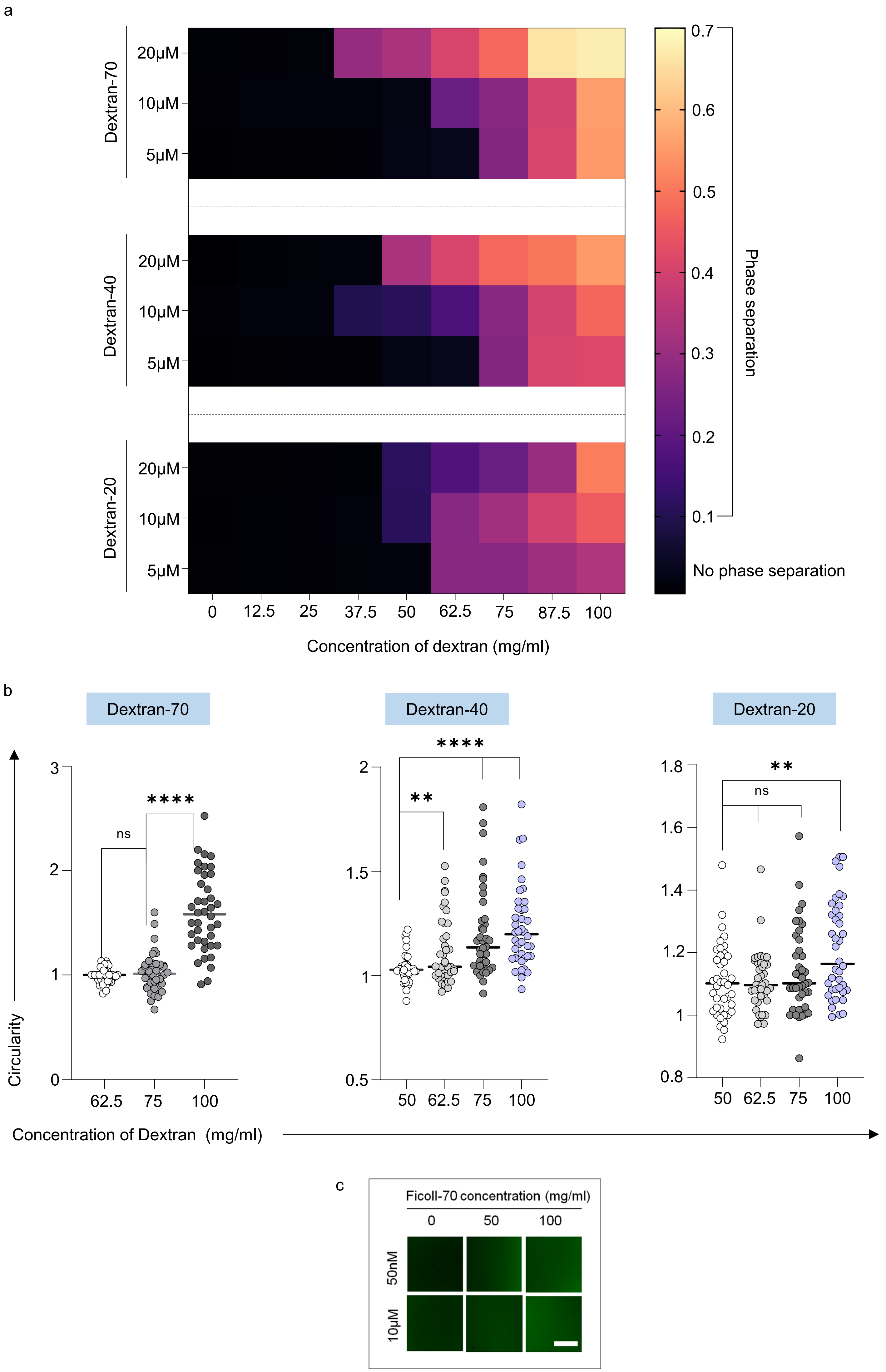

a
Dextran-70
Dextran-40
Dextran-20
20μM
10μM
5μM
0
12.5
25
37.5
50
62.5
75
87.5
100
Concentration of dextran (mg/ml)
0.7
0.6
0.5
0.4
0.3
0.2
0.1
Phase separation
No phase separation
b
Dextran-70
Dextran-40
Dextran-20
Circularity
Concentration of Dextran (mg/ml)
ns
****
**
ns
3
2
1
0
62.5
75
100
2
1.5
1
0.5
50
62.5
75
100
1.8
1.6
1.4
1.2
1
0.8
c
Ficoll-70 concentration (mg/ml)
0
50
100
50nM
10μM

**Figure S2:** (a) Turbidity plot of Sup35 in the presence of crowder: Turbidity was measured at 600nm. The protein concentrations are indicated in the left y-axis and dextran concentrations are indicated in the x-axis. The turbidity values are indicated in the colour bar. Below 0.1 there was no droplet formation confirmed by confocal microscopy. (b) Circularity plots of droplets at different Dextran concentrations in the phase separation regimes for different Dextran variants. Three independent measurements were taken with similar observations. Statistical significance was measured using paired t-test; ns: denotes nonsignificant; **P < 0.01; ****P < 0.0001 (c) Phase separation of Sup35NM in the presence of spherical crowder: Confocal microscopy images of 50 nM labeled Sup35NM in the presence of 50mg/ml and 100 mg/ml Ficoll-70(upper panel); Confocal microscopy images of 50 nM labeled Sup35NM mixed with 10μM unlabeled protein in the presence of 50mg/ml and 100 mg/ml Ficoll-70(lower panel). Scale bar: 5 μm.

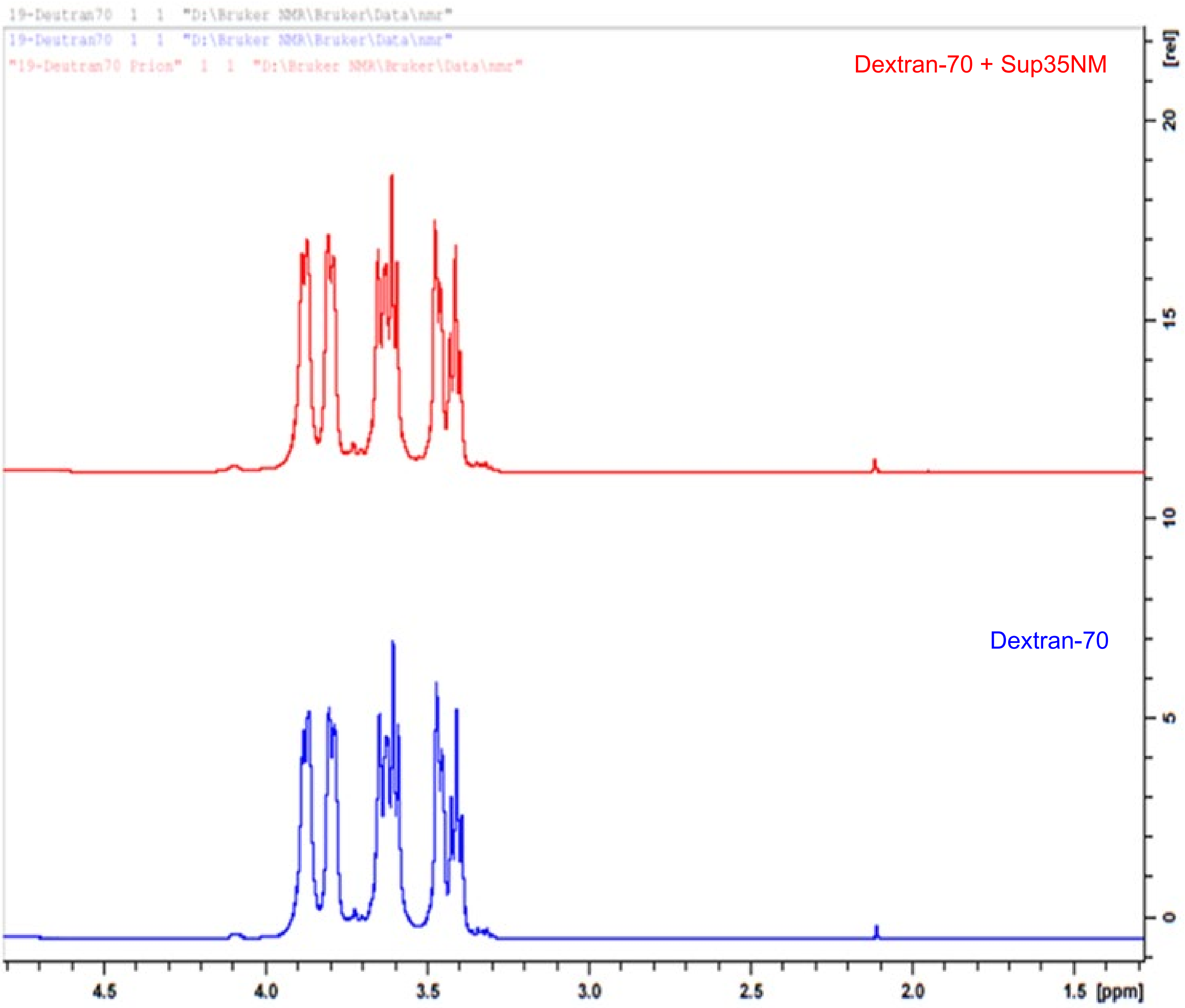

19-Deutran70 1 1 "D:\Bruker NMR\Bruker\Data\nmr"
19-Deutran70 1 1 "D:\Bruker NMR\Bruker\Data\nmr"
"19-Deutran70 Prion" 1 1 "D:\Bruker NMR\Bruker\Data\nmr"
Dextran-70 + Sup35NM
Dextran-70
[rel]
20
15
10
5
0
4.5
4.0
3.5
3.0
2.5
2.0
1.5
[ppm]

**Figure S3:** 1HNMR spectra of Dextran-70 in the presence and absence of Sup35. The red colour spectra represent the NMR spectra in the presence of Sup35NM and the blue colour spectra represent the NMR spectra in the absence of Sup35NM.

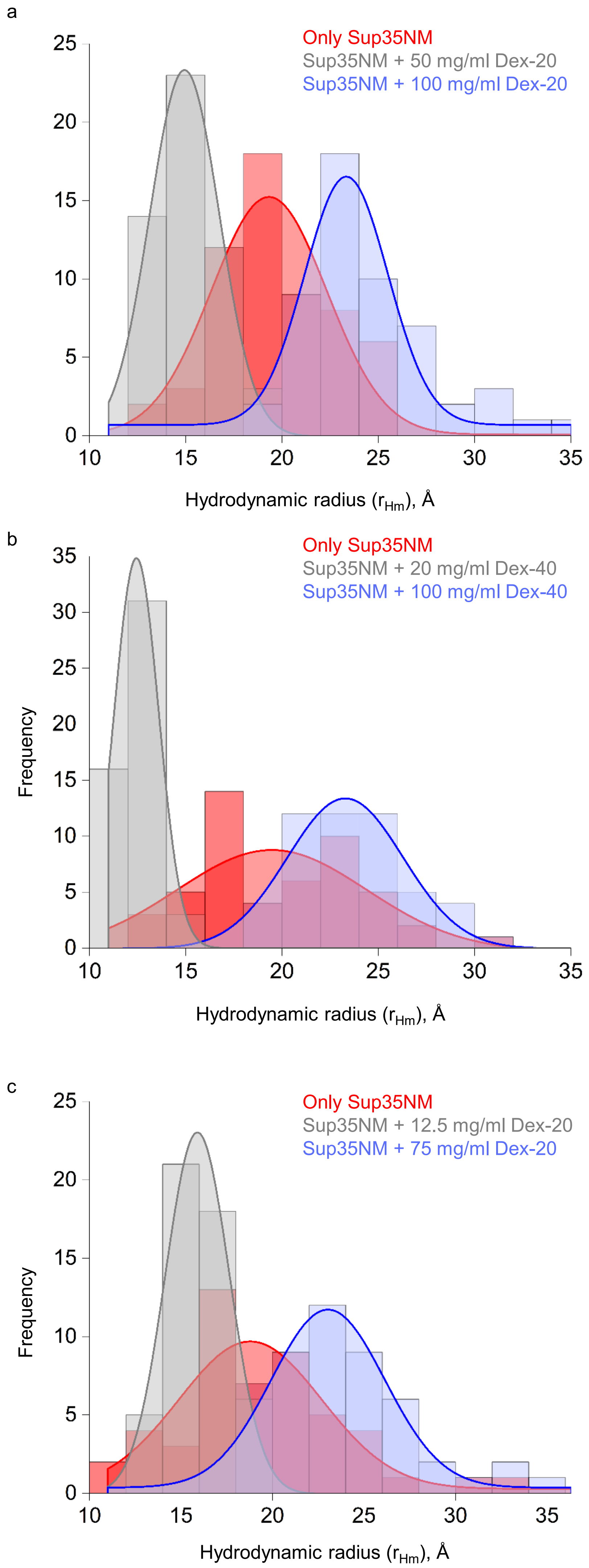

a
Only Sup35NM
Sup35NM + 50 mg/ml Dex-20
Sup35NM + 100 mg/ml Dex-20
25
20
15
10
5
0
10
15
20
25
30
35
Hydrodynamic radius ($r_{Hm}$), Å
b
Only Sup35NM
Sup35NM + 20 mg/ml Dex-40
Sup35NM + 100 mg/ml Dex-40
Frequency
35
30
25
20
15
10
5
0
10
15
20
25
30
35
Hydrodynamic radius ($r_{Hm}$), Å
c
Only Sup35NM
Sup35NM + 12.5 mg/ml Dex-20
Sup35NM + 75 mg/ml Dex-20
Frequency
25
20
15
10
5
0
10
15
20
25
30
35
Hydrodynamic radius ($r_{Hm}$), Å

Figure S4: Histogram plots of populations of Sup35NM under different types of Dextran. Concentrations of protein and Dextran are indicated in the respective figure. ~100 individual short FCS measurements were taken for each sample. (a) Histogram plots of hydrodynamic radius ($r_{Hm}$) of Native-like (red), Compact ( grey), and Extended (blue) populations of Sup35NM in the presence of Dextran-70. (b) Histogram plots of hydrodynamic radius ($r_{Hm}$) of Native-like (red), Compact (grey), and Extended (blue) populations of Sup35NM in the presence of Dextran-40. (c) Histogram plots of hydrodynamic radius ($r_{Hm}$) of Native-like (red), Compact ( grey), and Extended (blue) populations of Sup35NM in the presence of Dextran-20

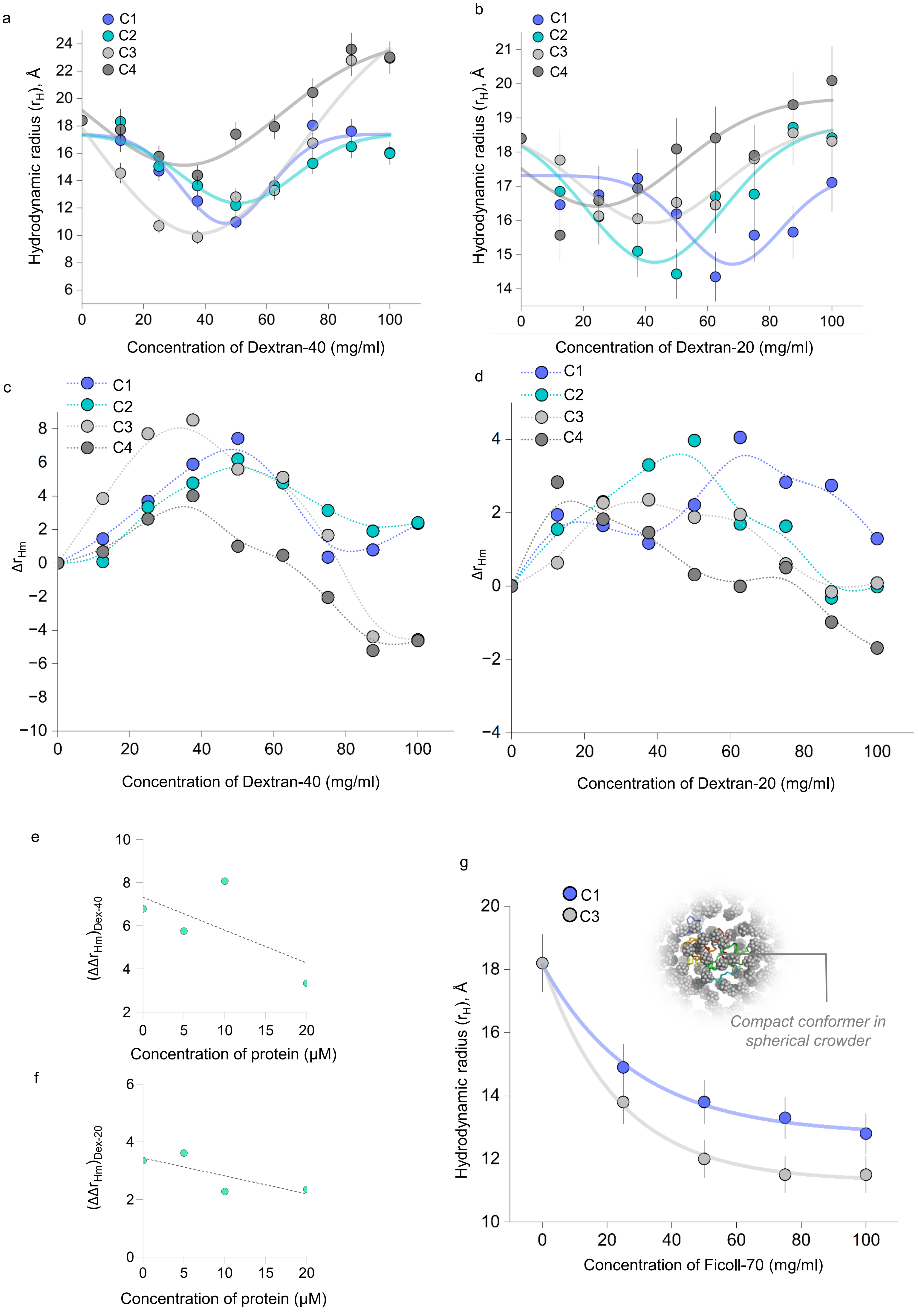
a
b
c
d
e
f
g
C1
C2
C3
C4
Hydrodynamic radius ($r_H$), Å
Concentration of Dextran-40 (mg/ml)
Concentration of Dextran-20 (mg/ml)
$\Delta r_{Hm}$
$(\Delta\Delta r_{Hm})_{Dex-40}$
$(\Delta\Delta r_{Hm})_{Dex-20}$
Concentration of protein (μM)
Concentration of Ficoll-70 (mg/ml)
Compact conformer in spherical crowder

**Figure S5:** (a) Plot of hydrodynamic radius ($r_{Hm}$), Angstrom vs. concentration (mg/ml) of crowder for Dextran-40. Purple circles indicate the 50nM Sup35NM, Cyan circles indicate 50nM labeled protein in 5μM unlabeled protein, Light grey circles indicate 50nM labeled protein in 10μM unlabeled protein, and Deep grey circles indicate 50nM labeled protein in 20μM unlabeled protein. All measurements were take n=3. The data are shown mean ± SD. (b) Plot of hydrodynamic radius ($r_{Hm}$), Angstrom vs. concentration (mg/ml) of crowder for Dextran-20. Purple circles indicate the 50nM Sup35NM, Cyan circles indicate 50nM labeled protein in 5μM unlabeled protein, Light grey circles indicate 50nM labeled protein in 10μM unlabeled protein, and Deep grey circles indicate 50nM labeled protein in 20μM unlabeled protein. All measurements were take n=3. The data are shown mean ± SD. (c) Plot of ($\Delta r_{Hm}$) at different crowder concentrations for Dextran-40 at varying protein concentration (C1-C4). (d) Plot of ($\Delta r_{Hm}$) at different crowder concentrations for Dextran-20 at varying protein concentration (C1-C4). (e) Plot of ($\Delta\Delta r_{Hm}$) vs. protein concentration indicated in the x-axis for Dextran-40. (f) Plot of ($\Delta\Delta r_{Hm}$) vs. protein concentration indicated in the x-axis for Dextran-20. (g) Plot of hydrodynamic radius ($r_{Hm}$), Angstrom vs. concentration (mg/ml) of crowder for Ficoll-70. Purple circles indicate the 50nM Sup35NM, Light grey circles indicate 50nM labeled protein in 10μM unlabeled protein. All measurements were take n=3. The data are shown mean ± SD.

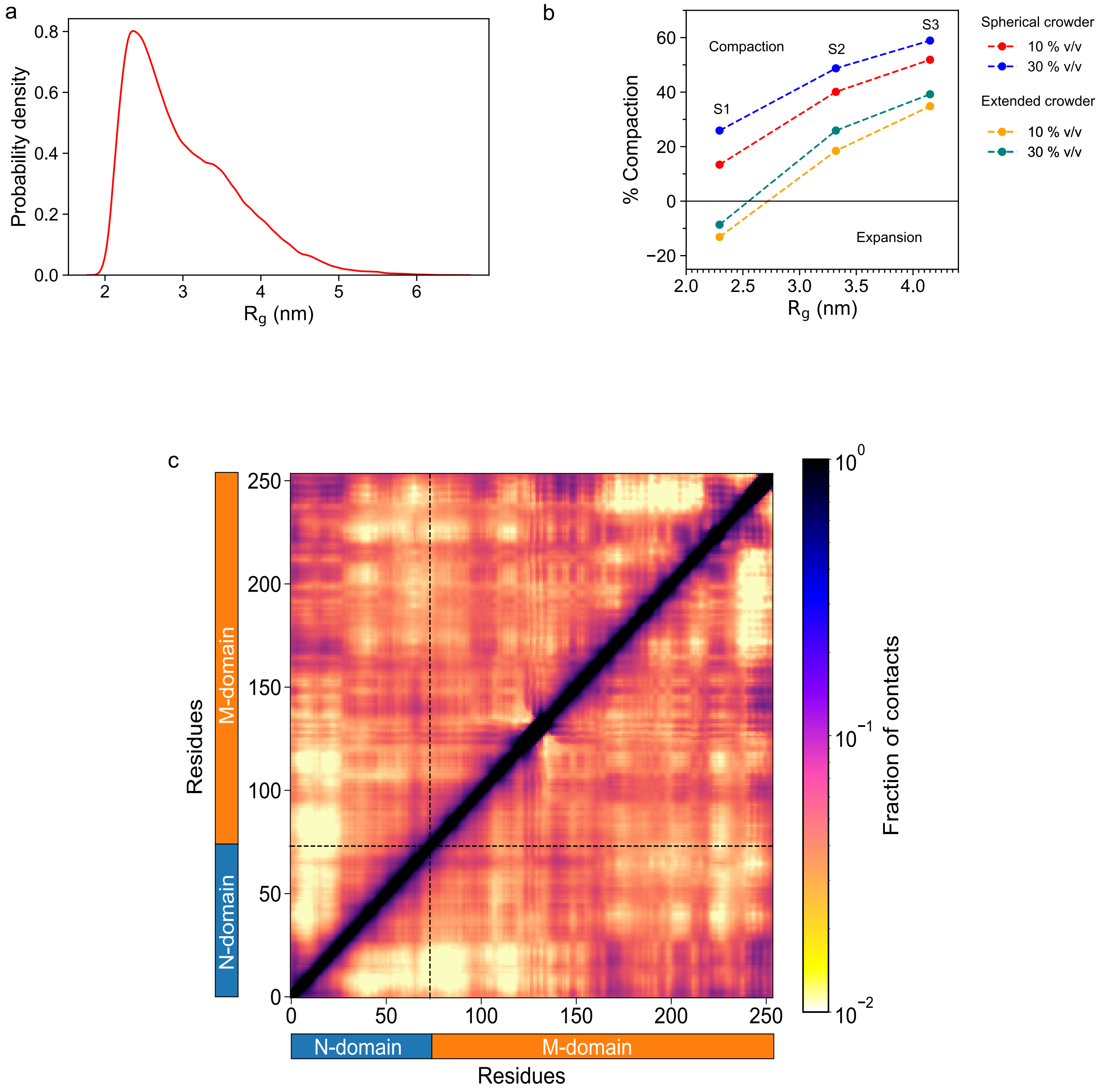
a
Probability density
0.0
0.2
0.4
0.6
0.8
2
3
4
5
6
$R_g$ (nm)
b
% Compaction
Compaction
Expansion
S1
S2
S3
60
40
20
0
−20
2.0
2.5
3.0
3.5
4.0
$R_g$ (nm)
Spherical crowder
10 % v/v
30 % v/v
Extended crowder
10 % v/v
30 % v/v
c
Residues
M-domain
N-domain
0
50
100
150
200
250
Fraction of contacts
$10^{0}$
$10^{-1}$
$10^{-2}$

**Figure S6:** CG simulation of Sup35NM: (a) Probability distribution of Rg of the ensemble. (b) Percentage compaction of three states (S1, S2 and S3) in the presence of 10 % v/v and 30 % v/v spherical and extended crowders. (c) Average intraprotein residue-wise contact probability maps of Sup35NM monomer in the presence of 30 % v/v spherical crowder. Axes denote the residue numbers. The color scale for the contact probability is shown at the extreme right of each plot of maps. The color bar along the axes of the plots represents the domains in the protein.

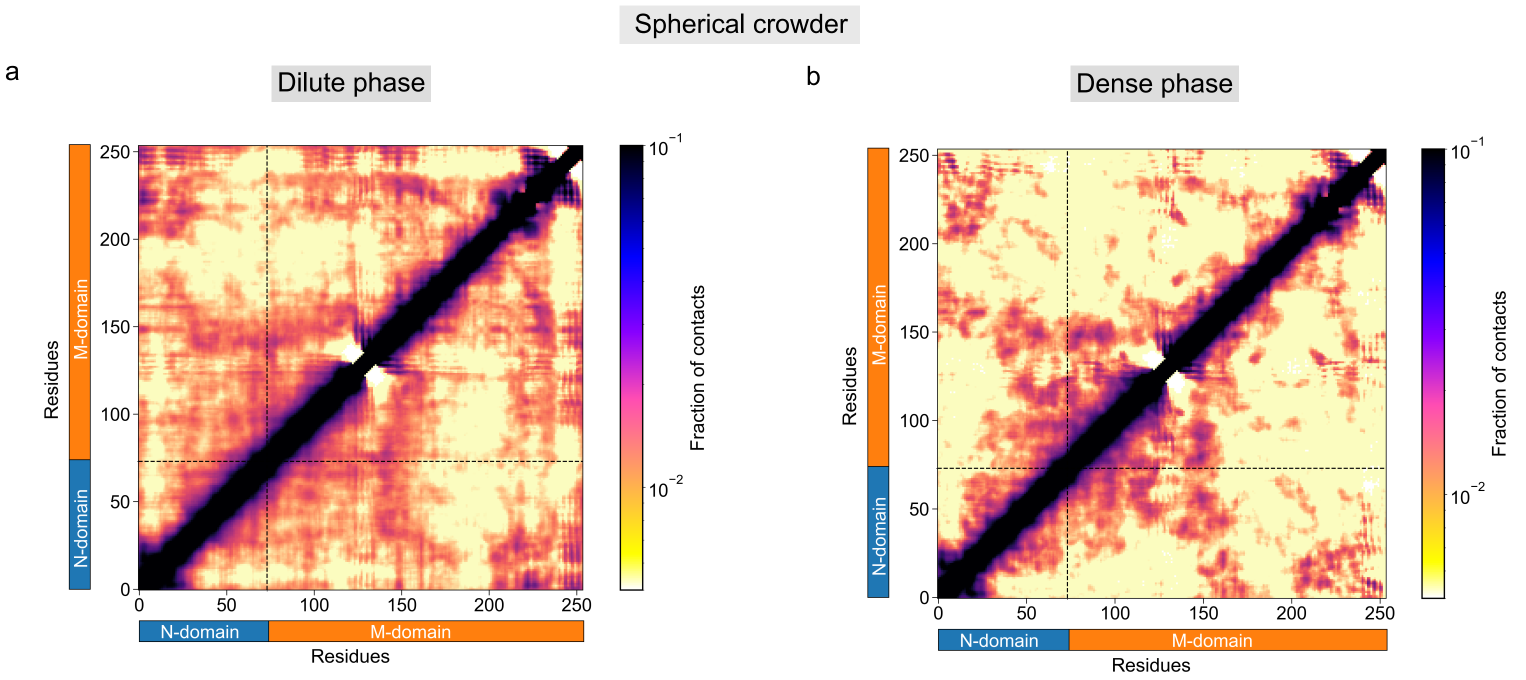

Spherical crowder
a
Dilute phase
b
Dense phase
Residues
N-domain
M-domain
Fraction of contacts
$10^{-1}$
$10^{-2}$
0
50
100
150
200
250

**Figure S7:** Average intraprotein residue-wise contact probability maps of Sup35NM in the (a) dilute and (b) dense (aggregate) phase in the presence of spherical crowder. Axes denote the residue numbers. The color scale for the contact probability is shown at the extreme right of each plot of maps. The color bar along the axes of the plots represents the domains in the protein.

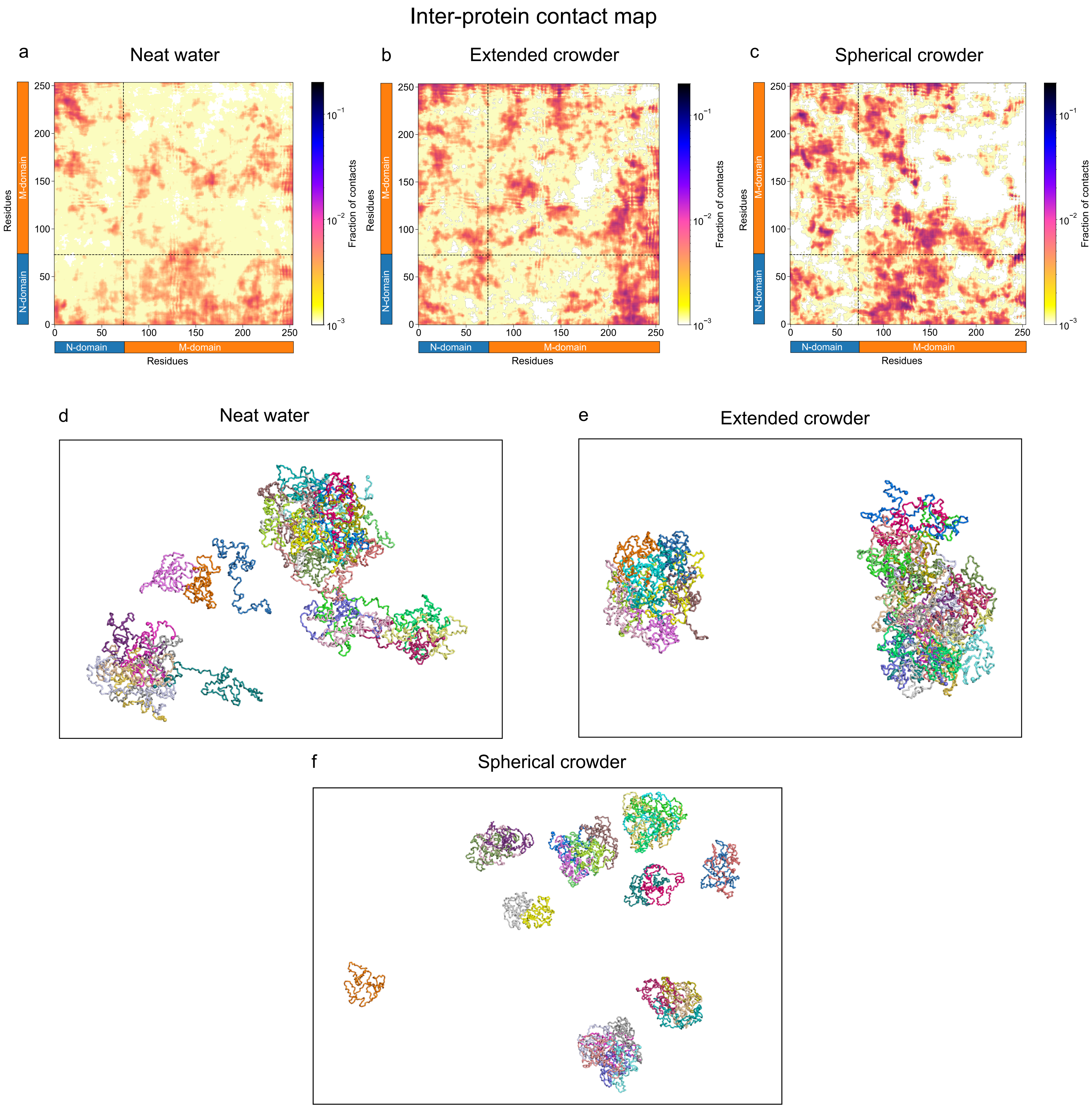

Inter-protein contact map
a
Neat water
b
Extended crowder
c
Spherical crowder
Residues
Fraction of contacts
N-domain
M-domain
0
50
100
150
200
250
$10^{-1}$
$10^{-2}$
$10^{-3}$
d
Neat water
e
Extended crowder
f
Spherical crowder

**Figure S8:** Average interprotein residue-wise contact probability maps of Sup35NM in aggregates formed in (a) neat water and in the presence of, (b) spherical crowder, (c) extended crowders. Axes denote the residue numbers. The color scale for the contact probability is shown at the extreme right of each plot of maps. The color scale for the contact probability is shown at the extreme right of each plot of maps. The color bar along the axes of the plots represents the domains in the protein. Snapshots of the aggregates formed in (d) neat water and in the presence of (e) extended crowders, (f) spherical crowder.
